# A RETINOBLASTOMA-RELATED transcription factor network governs egg cell differentiation and stress response in *Arabidopsis*

**DOI:** 10.1101/772400

**Authors:** Olga Kirioukhova-Johnston, Pallavi Pawar, Geetha Govind, Pramod Pantha, René Lemcke, Vidhyadhar Nandana, Danaé S. Larsen, Alagarsamy M. Rhahul, Jubin N. Shah, Patrick von Born, Chathura Wijesinghege, Yue Zhou, Wilhelm Gruissem, Franziska Turck, Maheshi Dassanayake, Amal J. Johnston

**Author notes:** Department of crop physiology, College of Agriculture, University of Agricultural Sciences, Hassan, India (Geetha Govind). Department of Plant and Environmental Sciences, University of Copenhagen, Frederiksberg, Denmark (René Lemcke). Division of Biochemistry, Freie Universitaet Berlin, Berlin, Germany (Vidhyadhar Nandana). Centre for Life Sciences, Peking University, Beijing, China (Patrick von Born, Yue Zhou). These authors contributed equally to this work (Pallavi Pawar, Geetha Govind).

## Abstract

The multicellular embryo, and ultimately the entire organism, is a derivative of the fertilized egg cell. Unlike in animals, transcription factor networks orchestrating faithful egg development are still largely unknown in plants. We have identified that egg cell differentiation in *Arabidopsis* require interplay between evolutionarily conserved onco-protein homologs RETINOBLASTOMA-RELATED (RBR) and redundant MYB proteins MYB64/MYB119. RBR physically interacts with the MYBs; and with plant-specific transcription factors belonging to the RWP-RK-domain (RKD) family and LEAFY COTYLEDON1 (LEC1), which participate in development of egg cells and inherent stress response. RBR binds to most of these egg cell-expressed loci at the DNA level, partially overlapping with sites of histone methylation H3K27me3. Since deregulation of *RKD*s phenocopies mutants of *RBR* and the *MYB*s in terms of cell proliferation in the egg cell spatial domain, all the corresponding proteins are likely required to restrict parthenogenetic cell divisions of the egg cells. Cross-talk among these transcription factors, and direct regulation by RBR, govern egg cell development and expression of egg-to-zygotic polarity factors of the WUSCHEL RELATED HOMEOBOX family. Together, a network of RBR-centric transcription factors underlies egg cell development and stress response, possibly, in combination with several other predicted nodes.

**Author summary:** The RETINOBLASTOMA protein is one of the core components of the Eukaryotic cell cycle, and corresponding evolutionary homologs have been implicated not only to repress cell division but also to control differentiation and development. How RETINOBLASTOMA RELATED (RBR) associate with other higher order regulators to control faithful egg cell development in sexual plants is pivotal for manipulation of successful reproduction in general, and engineering of parthenogenesis when asexual or apomictic seed progeny are desirable over sexual plants. Using a suite of molecular methods, we show that a RBR-associated transcription factor network operates to specify egg cells in *Arabidopsis*. Complex cross-regulation within these transcription factors seems to be necessary for successful maternal egg cell to zygotic transition and reproductive stress response. Detailed genetic analysis implicate that RBR and its interactive partners belonging to MYB and RWP-RK transcription factor families are possibly required to prevent parthenogenesis of the sexual egg cells. Novel RBR networks and stress nodes explained in this study might help to improve our understanding of sexual and asexual reproduction.

## Introduction

Proper differentiation of the egg cells is pivotal for sexual reproduction as well as parthenogenesis. In flowering plants, the egg cells are terminally differentiated within the miniature female gametophyte structures known as the embryo sacs that are encased by layers of sporophytic cells in the ovule. Cellular differentiation and maintained homeostasis are crucial for egg cell development, and they have been proposed to be orchestrated by positional cues during establishment of ovule and embryo sac polarity [1–6], and ultimately of the egg cell and zygote [7, 8] in *Arabidopsis*. Tightly coordinated developmental processes implicate both directed cell-to-cell communication and cell-autonomous regulation operating throughout embryo sac development and fertilization processes.

Egg cell development in plants is proposed to be under the control of molecular factors including cell cycle regulators, transcription factors, RNA splicing machinery, signalling molecules such as secreted peptides and chromatin dynamics {reviewed in [1]}. Transcription factors play a predominant role in regulation of gene expression, thus, tight control over transcriptional regulation is foreseeable in the egg cell [9, 10]. A microarray expression analysis of the *Arabidopsis* egg cell transcriptome suggests that >350 transcription factors could be expressed there [9]. Although this is likely an underestimate, considering the difficulty in isolating the *Arabidopsis* egg cell over that of rice [10], large transcription factor families such as MYB, RWP-RK domain-containing (RKD) and WUSCHEL-RELATED HOMEOBOX (WOX) have been proposed to be prominent members of the egg cell transcriptome [1, 9]. Functional dissection of these egg cell-expressed transcription factors, and exploring the inherent cross-talks and associated networks, will give a clear picture of egg cell determination and patterning in plants.

A unique feature of the egg cell in sexually reproducing organisms is a temporary arrest of its cell divisions until fertilization and reprogramming to zygotic gene expression. In flowering plants, deregulation of a homologue of *BABYBOOM (BBM)* [11], *MULTI-SUPPRESSOR OF IRA 1 (MSI1)* [12] and specific R2R3-type *MYBs* [13] are implicated in autonomous developmental events in the embryo sac and in particular the egg cell. In *Arabidopsis*, overexpression of *WUSCHEL*, *MYB* genes, *BBM*, and *LEAFY COTYLEDON1 (LEC1)* has been shown to induce somatic embryogenesis [14–16], indicating that they participate in transcriptional rewiring towards embryo development. *LEC1* encodes a CCAAT-box binding transcription factor known primarily for its role during embryogenesis, seed maturation and stress amelioration [17, 18]. Interplay of WUSCHEL-related transcription factors WOX2 and WOX8 prepares the egg cell to establish zygote polarity [8]. Unlike the neighbouring central cell, the egg cell chromatin environment is rather transcriptionally quiescent with high levels of repressive histone methylation marks such as H3K27me3 [19, 20]. Nevertheless, combinatorial transcription factor regulation and epigenetic modifications likely play an important role during egg cell development in plants.

Ready for fertilization, the egg cell in *Arabidopsis* likely stays quiescent in the G2-phase of the cell cycle, thus matching the cell cycle stage of the sperm cell at the onset of fertilization [21]. Only a few cell cycle regulators are expressed in the egg cell [9], including the higher-order transcriptional repressor RETINOBLASTOMA RELATED (RBR). Deregulation of RBR has been shown to perturb egg cell specification and genome integrity [22–24]. While RBR and its paralogues control transcriptional networks in somatic cell types, either dependent or independent of the cell cycle [25, 26], whether they play a similar role in egg cell development is currently not understood. In this study, we have established functional links between RBR and a subset of transcription factors controlling egg cell development and stress response during *Arabidopsis* reproduction. Notably, we have demonstrated the importance of an RBR-centric egg cell-expressed transcription factor network essential for sexual and parthenogenetic reproduction, and have identified several putative nodes of this network for further dissection.

## Results

### *RBR* is required for egg cell development

In order to understand the true expression of RBR during egg cell development, we constructed a reporter line consisting of an N-terminal fusion of GFP to a genomic *RBR* locus that was driven under its 2.2 Kbp promoter and included 3’ flanking region (*pRBR*::*GFP-RBR*). To test the functionality of the construct, first we introduced the transgene into the amorphic *rbr-3 Arabidopsis* mutant in which the entire embryo sac is defective and cells proliferate instead of appropriate cell differentiation [6, 24]. Screening multiple independent transformants, we recovered *pRBR*::*GFP-RBR* lines that were able to completely restore the wild-type function of *RBR* in the female gametophytes. These transgenics fully rescued the *rbr-3-* mediated ovule sterility, as evident from restored seed set, and transmission of the mutant allele to the progeny (Fig 1A-B, Table S1-S2). Therefore, the *pRBR*::*GFP-RBR* was sufficient for development of the embryo sac including the egg cell. Next, we examined these transgenics for expression of GFP, which would correspond to the endogenous localization of RBR. Consistent with the phenotypic complementation, we detected the recombinant protein produced by the *pRBR*::*GFP-RBR* construct in the synergids and the egg cell (Fig 1C). Upon egg cell fertilization, the *pRBR*::*GFP-RBR* signal decreased in the zygote to almost undetectable level (Fig 1C-E). Thus the egg apparatus expression of RBR under its native promoter was sufficient for proper egg differentiation, and a rapid reduction of RBR upon fertilization must have been important for egg-to-zygote transition.

**Fig. 1.**
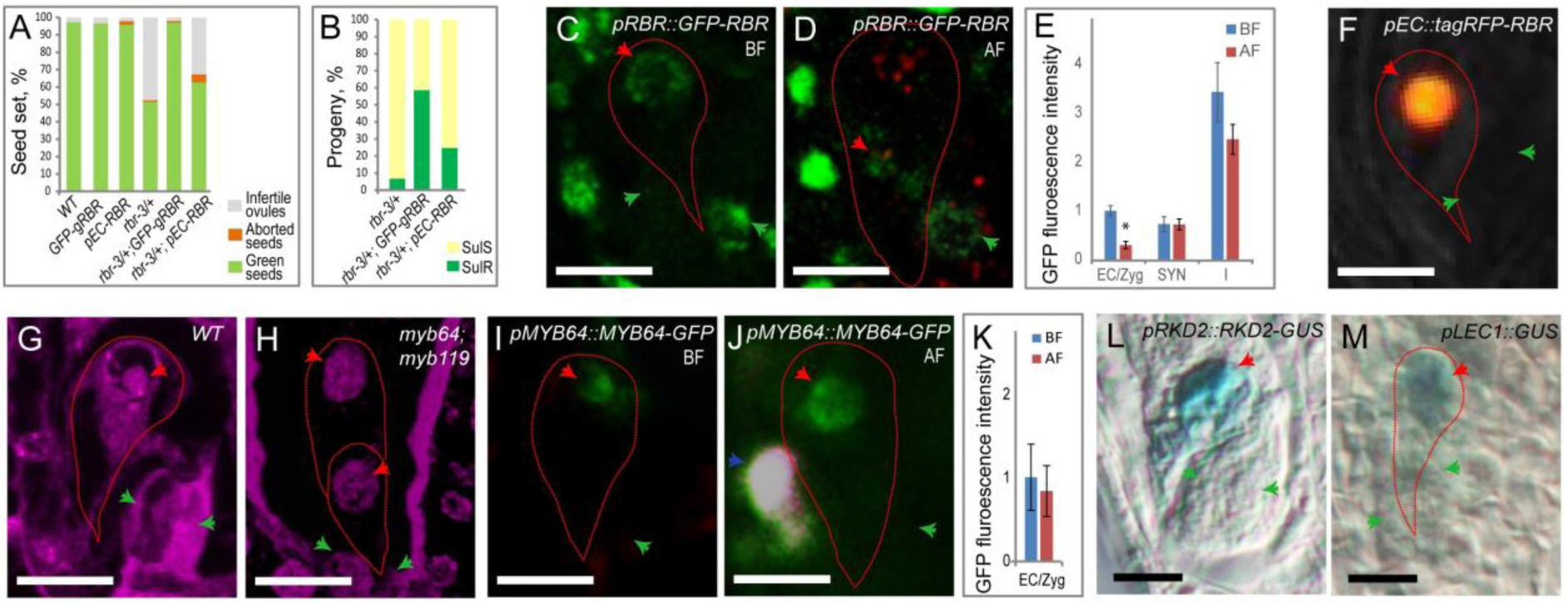
RBR-MYB transcription factors are essential for faithful egg cell development. (**A**) Rescue of ovule/seed abortion in *rbr-3* plants in presence of the two RBR constructs. (**B**) Transmission of *rbr-3* allele to the progeny in presence of three tagged RBR constructs, scored by resistance to Sulfadiazine (Sul, S - sensitive; R-resistant). (**C**) eGFP fused to genomic RBR is detectable in the mature egg cell. (**D**) GFP-gRBR signal is largely depleted in the polarized zygote upon fertilization (∼8 hours after pollination, hap). (**E**) Quantification of GFP-RBR signals before and after fertilization (BF, AF). (**F**) Egg cell-specific tagging of RBR by *tagRFP-RBR* fusion. (**G-H**) Feulgen-stained female gametophytes: (**G**) A mature wild-type (WT) embryo sac showing an egg and other cell types such as two synergids and a central cell. (**H**) Loss of *MYB64/MYB119* leads to embryo sac proliferation. (**I**) MYB64-GFP protein in the mature egg cell. (**J**) MYB64-GFP upon pollen tube entry (∼8 hap). (**K**) MYB64-GFP signals quantified. (**L**) RKD2-GUS translational fusion is localized to the egg cell only. (**M**) *pLEC1*-GUS is faintly expressed in the mature egg cell-containing embryo sac. Red/green/blue/white arrow-heads: egg/synergids/sperm/central cell. Scale bar=20µm.

To test whether RBR expression in the egg cell alone is sufficient to restore its wild-type function, we expressed a *tagRFP-RBR* fusion under a strong egg cell-specific promoter of *EGG CELL 1.1* (*pEC)* [27] (*pEC::tagRFP-RBR*) (Fig. 1F) and introduced it into the *rbr-3* mutant too. Indeed, *pEC::tagRFP-RBR* construct could partially rescue the null *RBR* mutation as evident from improved seed set and strong increase of *rbr-3* allele transmission to the progeny from 7% in *rbr-3/+* alone [22] to 25% in the presence of the transgene (Fig.1A-B; Table S3-S4). Notably, the strong overexpression of *RBR* in the egg cell, as visualized by expression of tagRFP, did not cause obvious aberrations in the embryo sac nor seed development in the wild-type background. Taken together, data as above suggest that the amount of RBR under its native promoter is sufficient for egg cell development, while its increased dosage in the egg cell does not perturb sexual reproduction.

### RBR-dependent transcriptional regulation in the egg cell

Since abolishing RBR expression caused severe perturbations in egg cell differentiation, development and function [22], and that these developmental anomalies could be restored when RBR is expressed specifically in egg cell (Fig. 1A-B,F), we reasoned that most of the phenotypic effects observed in the *rbr-3* mutant were due the absence of RBR expression in the egg cell. Therefore, we made use of the amorphic *rbr-3* mutant to uncover the transcriptional changes in the egg cell upon depletion of *RBR* by comparing transcriptional profiles of the mutant ovules against the wild-type ovules. We assumed that differences between transcriptomes of *rbr-3* and wild-type ovules should reflect mainly the female gametophyte-specific gene expression [2,6,28], for *RBR* gene is haplosufficient both in sporophytic and gametophytic tissues [22]. We conducted a two-step comparative transcriptome analysis. In step-1, all the acquired mRNA-seq data from the ovules were filtered to retain those genes reported as part of the *Arabidopsis* egg cell microarray dataset [9]. In step-2, the filtered data were subjected for wild-type versus mutant differential expression analysis. This approach allowed us to scrutinize egg cell-related gene expression profiles of the wild-type versus *rbr-3* genotypes.

Differential gene expression analysis identified a total of 2096 egg cell-expressed genes with over 20 transcripts that were previously validated for spatial egg cell expression (Table S5, Fig. 2A-B). GO enrichment analysis of the egg cell-expressed transcripts pinpointed several transcription factors (105) and stress-related genes (172) that potentially could function downstream of *RBR* (Table S6, Fig. S1). Among *rbr-3* up- and down-regulated genes, 120 and 70 candidates, respectively, (ca. 9%) fell under GO categories related to stress. Interestingly, 18% of transcription factors upregulated in *rbr-3* were stress-related versus 7% that were down-regulated, indicating a predominant repressive function of RBR on transcriptional regulators involved in stress responses in the egg cell.

**Fig. 2.**
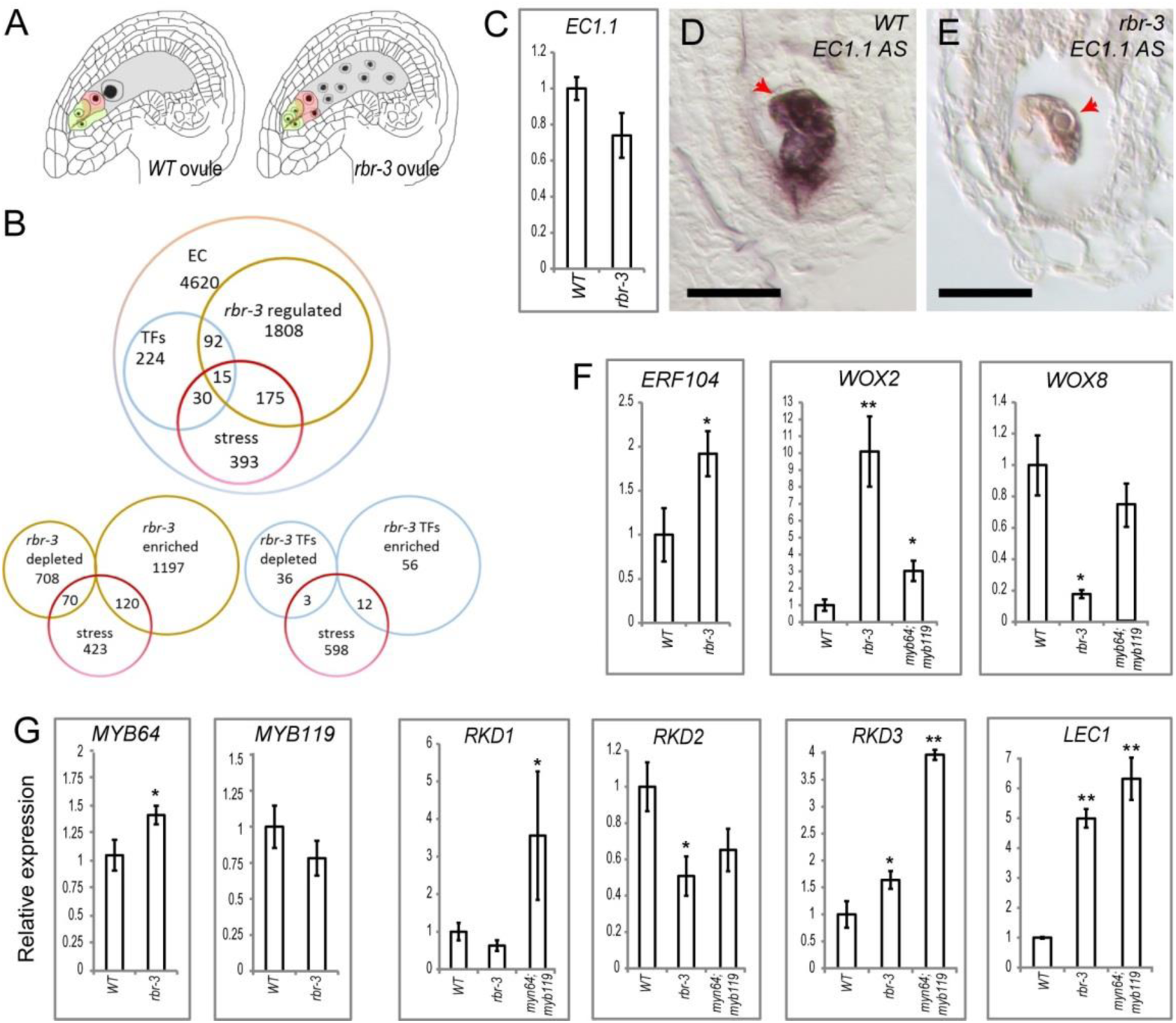
Abrogation of RBR causes global egg cell-expressed gene deregulation. (**A**) A schematic of ovule samples used for differential RNA-seq analysis. (**B**) Genetic subtraction of egg cell-expressed genes from the ovule transcripts identifies genes regulated by RBR in the egg cells. (**C,F,G**) Validation of RBR-regulated candidate genes by real-time qRT-PCR in *rbr-3*, and testing in *myb64;myb119* ovules. Significance **α ≤ 0.01; *α ≤ 0.05. (**D-E**) Spatial validation of RBR-regulated *EC1.1* by mRNA *in situ* hybridization.

In order to validate the mRNA-seq data and to build up a functional egg cell-associated transcriptional regulatory network, we chose specific differentially expressed candidate genes that have been known for their egg cell expression [9] or function. We confirmed downregulation of egg cell-expressed genes *EGG CELL 1.1* (*EC1.1*) [27] and *WOX8*, and upregulation of *WOX2* [8, 29] and *ETHYLENE RESPONSE FACTOR 104 (ERF104)* [9] by qRT-PCR (Fig. 2C-F). In addition, abundant *EC1.1* mRNA *in situ* signals were readily visible in the wild-type egg cell; however, in *rbr-3* eggs, *EC1.1* signals were partially depleted (Fig. 2D-E). Validating the spatio-temporal gene expression profile of egg-cell specific markers such as *EC1.1* served as a quality control of our genetic subtraction approach.

While the egg-like cells of *rbr-2* embryo sacs still expressed promoter reporters of female gametophyte-expressed MYB transcription factors *pMYB64::GFP* and *pMYB119::GFP* [13], our mRNA-seq analysis on *rbr-3* ovules identified *MYB64* as a differentially expressed transcript (Table S5). Indeed, validation by real-time qRT-PCR confirmed a slight but statistically significant upregulation of *MYB64* in *rbr-3* ovules (Fig. 2G). MYB64 was found to be functionally redundant with MYB119, and double *myb64;myb119* mutants showed severe aberrations in embryo sac development including egg-cell like proliferation almost phenocopying amorphic *rbr* mutations [6, 22] (Fig. 1G-H). It is interesting to note that loss of *MYB64;MYB119* function had a similar effect on *WOX2* and *WOX8* expression (Fig. 2F). Several double allelic combinations of *myb64* and *myb119* showing similar embryo sac proliferation phenotypes were elaborated, and were fully rescued in presence of intact *MYB64* or *MYB119* [13]. We detected mRNA for both genes throughout the mature embryo sac (Fig. S2), confirming that the previously analysed promoter::GFP fusions [13] reflected endogenous expression patterns of the corresponding loci. Further, we noticed that, unlike pRBR::GFP-RBR, pMYB64::MYB64-GFP protein was rather abundant in the unfertilized egg and in the early zygote (Fig. 1I-K). The transcriptional profile of most of the *RKD* genes pinpointed their preferential expression in the *Arabidopsis* egg cell [9, 30]; however no reports are available on the corresponding proteins. We found that a RKD2 protein fusion with GUS (β-glucuronidase) was localised to the egg cell in *Arabidopsis* (Fig. 1I). *RKD2* was significantly downregulated at least in *rbr-3* (Fig. 2G). Although *RKD1* gene expression was slightly down-regulated in *rbr-3* ovules, it was upregulated in *myb64;myb119* mutant. In contrast, *RKD3* was upregulated in both mutants (Fig. 2G). We found that LEC1, a stress-related gene primarily expressed during seed development, is also expressed in the egg cell. Expression of *pLEC1::GUS* construct was undetectable in the sporophytic cells of the ovule; however, a faint GUS signal was recorded throughout the embryo sac including the egg cell (Fig. 1L). Similar to *RKD3, LEC1* transcripts were strongly upregulated in *rbr-3* and *myb64;myb119* ovules (Fig. 2G). Since *RKD2*, *RKD3* and *LEC1* are commonly deregulated in both *rbr-3* and *myb64;119* mutant ovules that phenocopy each other, it is apparent that RBR and MYB64/119 act in the same regulatory pathway upstream of RKDs and LEC1 transcription factors during egg cell development.

### RBR is tethered to promoters of a subset of egg cell transcription factors

To establish whether the egg cell genes upregulated in the *rbr-3* mutant could be direct targets of RBR, we performed chromatin immunoprecipitation (ChIP) experiments. We tested enrichment of promoter fragments of *MYB64*, *LEC1*, *RKD3* and *WOX2* after immunoprecipitation of GFP-RBR in reproductive tissues that contained mature egg cells (Fig. 3A-B). A fragment spanning an E2F binding site in promoter of *PROLIFERATING CELL NUCLEAR ANTIGEN* (*PCNA*)served as a positive control for RBR binding, and promoter of *At1g69770* as a negative control [31]. Upon antibody background subtraction, most of the tested fragments were found RBR-associated (Fig. 3B-C). *pRKD3, pWOX2* and *pLEC1* fragments were enriched in both RBR-ChIPs across vegetative and reproductive stages (Fig. S3A). The *MYB64* promoter, however, showed differential RBR occupation level at two tested E2F binding sites between the tissue types. Surprisingly, the *MYB64* gene body was also bound by RBR. In order to verify RBR targeting both MYB64 and RKD3 directly in the egg cell, we performed an additional ChIP experiment using *pEC::tagRFP-RBR* transgenics. Here, we examined specific sites of *pMYB64* (containing predicted canonical E2F sites) and *pRKD3* (no E2F sites) promoters that we tested earlier for binding by GFP-RBR, upon immuno-precipitation of tagRFP-RBR. Both *pMYB64* and *pRKD3* promoter fragments were found enriched for this epitope binding (Fig. 3C), leading us to conclude that RBR represses transcription of *MYB64* and *RKD3* in the egg cell by directly binding to their corresponding promoters.

**Fig. 3.**
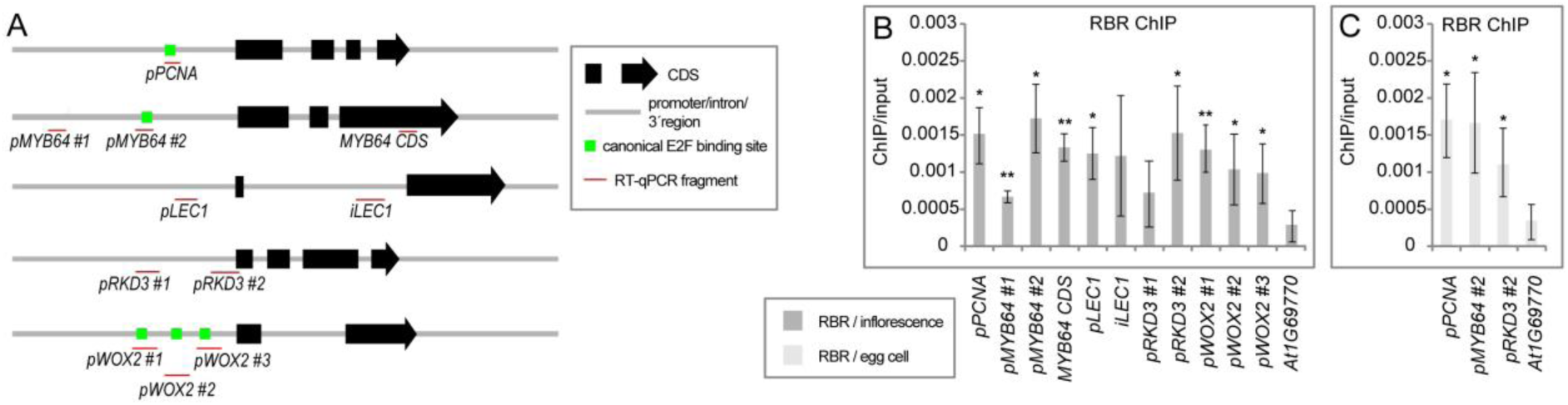
RBR associates with gene promoters of MYB and RKD families of transcription factors. (**A**) Location of RBR protein interaction with DNA fragments tested in Chromatin-Immunoprecipitation for RBR or for the PRC2-specific H3K27me3. (**B-C**) ChIP in reproductive tissues containing mature egg cells. Relative ChIP-qPCR normalized by input. Significant difference between the negative control (background) and the experimental values: **α ≤ 0.01; *α ≤ 0.05.

Next, we asked if the sites of RBR binding within our target egg cell candidate loci overlapped with the repressive histone methylation mark H3K27me3, similar to previous findings in seedling tissues [31]. We performed ChIP for H3K27me3-bound DNA in egg cell-containing gynoecia using a ChIP experiment on seedlings as a baseline for chromatin occupation by H3K27me3 (Fig. S3B). A previously reported fragment with an E2F binding site in the *PCNA* promoter was used as a negative control for H3K27me3 enrichment [31]. H3K27me3 mark loading in gynoecia tissues resembled overall that of the seedlings, with significantly lower and higher enrichment in *RKD3* and *WOX2* promoters, respectively. In both the seedlings and reproductive tissues, *MYB64* showed low H3K27me3 binding in its promoter similar to *pPCNA*, and high binding at the coding regions. The *LEC1* locus was moderately decorated with H3K27me3 mark. Our data pinpointed that both RBR and the Polycomb Repressive Complex 2 (PRC2) with its inherent H3K27me3 activity could bind to the promoters of the egg cell-expressed transcription factors investigated.

### Deregulation of RETINOBLASTOMA network partly phenocopies stress-induced effects on egg cell development

Both *rbr-3* and *myb64;myb119* female gametophytes mostly fail to arrest mitotic divisions, often showing multiple cells at the position of the egg cell and other cell types (Fig. 1G-H, 2A, 4A-C). In *Arabidopsis,* there are five *RKD* genes, but their function during egg cell development is masked due to redundancy and lack of faithful mutant alleles [30, 32]. We previously reported that three *RKD* genes are preferentially expressed in the egg cell, and act as activators of a subset of unknown genes expressed there [30] (Fig. 1L). Two other RKDs are also expressed in the egg cells and also elsewhere in the sporophyte [9, 33]. Therefore, we analysed the role of the RKD family by attaching a transcriptionally repressive EAR-domain [34] to RKD2 driven by the egg cell-specific *pEC* promoter (referred to as *pEC::RKD^DN^)*. Stable expression of the *pEC::RKD^DN^* transgene in plants led to variable seed set ranging from 50-75% of viable seeds, and the remaining ovules aborted at early stages. A number of mutant embryo sacs had additional egg-like cells in the egg apparatus (N=14/123) (Fig. 4D-E), and some embryo sacs completely collapsed (Fig. 4F). Most strikingly, when fertilization was blocked, we observed rare cases (N=5/81) of parthenogenetic zygote/embryo development that subsequently aborted as the unfertilized central cell failed to produce the endosperm (Fig. 4H-I, compare to sexual zygote in 4G). Taken together, deregulation of the RKD factors partially resembled loss of RBR or MYB64;MYB119 activity in terms of additional egg cells within the same embryo sac, supporting that they are down-stream and/or a part of the RBR and MYB pathway operating in the egg cell specification.

**Fig. 4.**
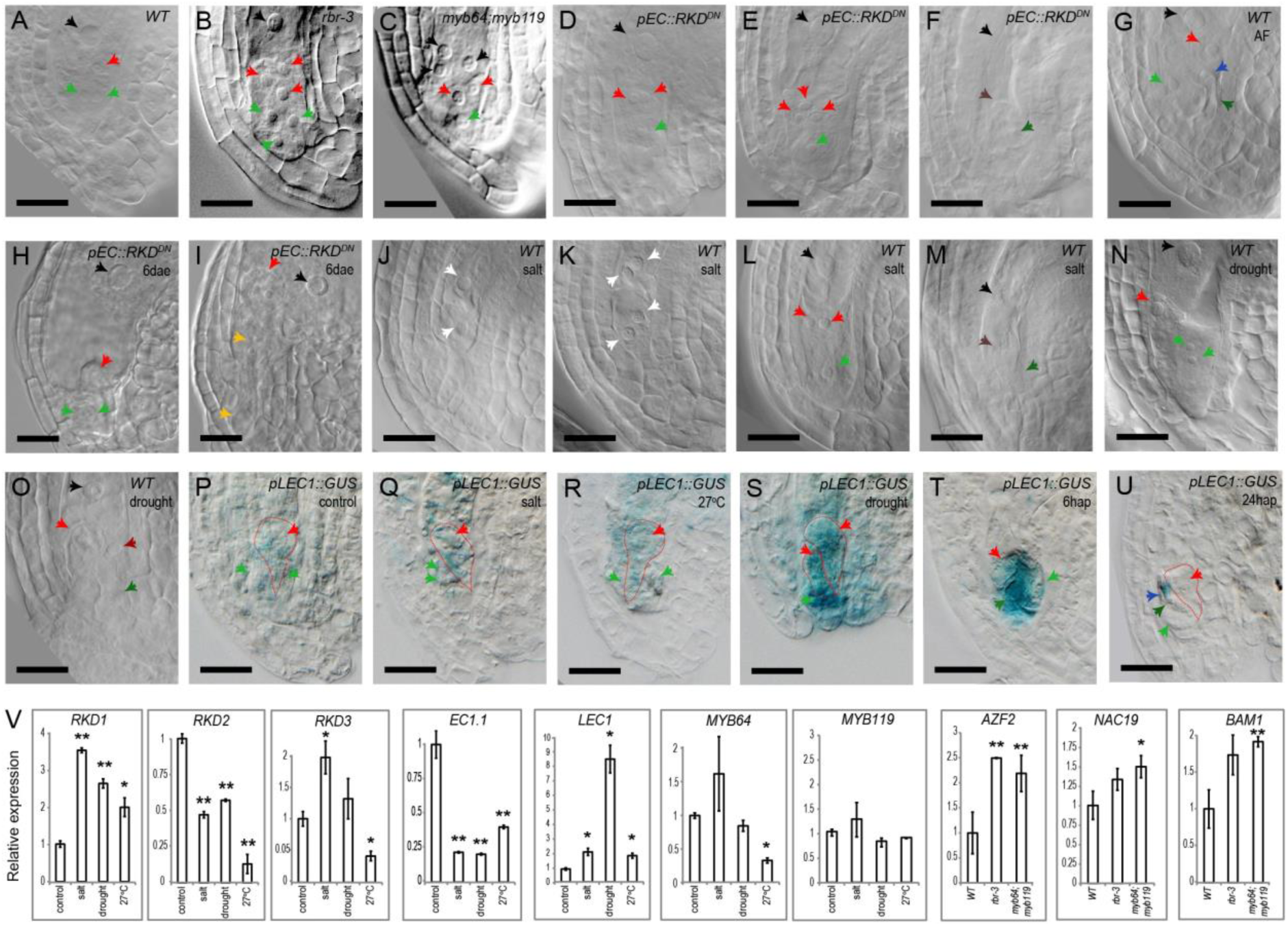
Abiotic stress and deregulation of RBR network leads to additional egg cells. (**A**) A WT embryo sac with mature egg cell. (**B-D**) Supernumerary eggs in *rbr-3* and *myb64;myb119*, and *pEC::RKD^DN^* embryo sacs. (**E**) A *pEC::RKD^DN^* ovule with rare three egg cell-like phenotype or completely collapsed (**F**). (**G**) A sexual zygote in WT at ∼12 hap. (**H-I**) rare fertilization-independent zygote-like structure or parthenogenetic embryo in *pEC::RKD^DN^* ovules 6 days after emasculation. (**J-M**) salt-stress induced embryo sac defects and twin eggs. (**N**) WT-looking drought-treated ovule. (**P-U**) *LEC1* expression in the embryo sac under salt stress (Q), elevated temperature (27°C) (**R**), and drought (**S**). *LEC1* is activated in the synergid upon pollen tube entry ∼ 6 hap (**T**), and only residual *LEC1* is detectable when the zygote is polarized a day after pollination (**U**). Scale bar = 20µm. hap – hours after pollination. See Fig. 1 for color-scheme of arrow-heads, brown – collapsing embryo sac; yellow – suspensor. (**V**) Relative expression in mature egg cell-containing gynoecia under stress. Significance **α ≤ 0.01; *α ≤ 0.05

Several stress genes were deregulated in *rbr-3* grown in ambient conditions (Fig. 2A-B, 4V, S1), suggesting activation of a cellular stress response in the absence of RBR function in the embryo sac. We noted that egg cell-expressed stress-response genes such as *ARABIDOPSIS ZINC FINGER 2 (AZF2), NAC19, and BETA-AMYLASE 1 (BAM1)* were upregulated not only in *rbr-3* but also in *myb64;myb119* ovules (Fig. 4V). We tested our hypothesis of stress influencing embryo sac or egg cell development by exposing soil-grown plants to external abiotic stress conditions such as NaCl, drought, and elevated temperature (27°C). Under salinity stress, we observed desynchronization of embryo sac development ranging from two-nucleate to mature four-celled stages in the same flower, while in control 95% of ovules contained mature embryo sacs (n=157 and 174), indicative of delayed egg cell development under salt stress. Furthermore, we observed formation of twin egg cells, and also collapsed embryo sacs under salinity (Fig. 4J-M), while mild drought led to rather wild-type like ovules with rare observations of additional egg-like cell along with the fertilized zygote (Fig. 4N-O).

The *LEC1* reporter was upregulated in the egg cells upon different stress conditions (Fig. 4P-S,V), substantiating the anticipated role of LEC1 in mediating stress response in the egg cell similarly to other plant organs [reviewed in [35]]. Additionally, very strong activation of *LEC1* gene was observed in the egg apparatus specifically in synergids upon pollen tube entry, suggesting a strong stress response during programmed cell death of synergid cells (Fig. 4T-U). *RKD* expression responded to stress in a distinct manner. Both *RKD1* and *RKD3* were mainly upregulated and *RKD2* was downregulated upon stress, suggesting differential regulation across these redundant and recently-duplicated factors. *MYB64* and *MYB119* expression was not significantly altered across most stress conditions, except for downregulation of *MYB64* under elevated temperature. Whether it is (a) abiotic stress or (b) mutational effects in *rbr-3* and *myb64;myb119* grown under ambient conditions, it is noteworthy that phenotypic effects like induction of super-numerary egg cells and transcriptional responses for RBR-regulated genes, are similar across experiments.

### Expanding the RBR-centric egg cell network

In addition to the transcriptional regulation centred on RBR and the egg cells in *Arabidopsis*, we asked if other regulatory cues such as protein-protein interaction can also be identified for the egg cell-expressed proteins. First, we performed a heterologous two-hybrid protein-protein interaction experiment in yeast, using RBR as bait and the egg cell-expressed transcription factors as prey. We used an RBR-interacting protein MULTI-SUPRESSOR-OF-IRA 1 (MSI1) as a prey in control experiments [28]. When RBR was used in binary combinations with MSI1, MYB64, MYB119, RKD1, RKD2, RKD3 and LEC1, we observed growth of yeast cells in appropriate drop-out media, hinting that interaction between RBR and these transcription factors occurred heterologously in yeast (Fig. S4). We validated these interactions *in vivo* in plant cells by using Bi-molecular Fluorescence Complementation (BiFC). The transient BiFC in tobacco leaves confirmed that RBR interacted with MYB64 and MYB119, RKD1-3 and LEC1, although the signals were rather weak in case of MYB119 and LEC1 (Fig. 5A-B). Concisely, we identified three different groups of transcription factors as part of the RBR egg cell regulatory network, among which two represent redundant gene families.

**Fig. 5.**
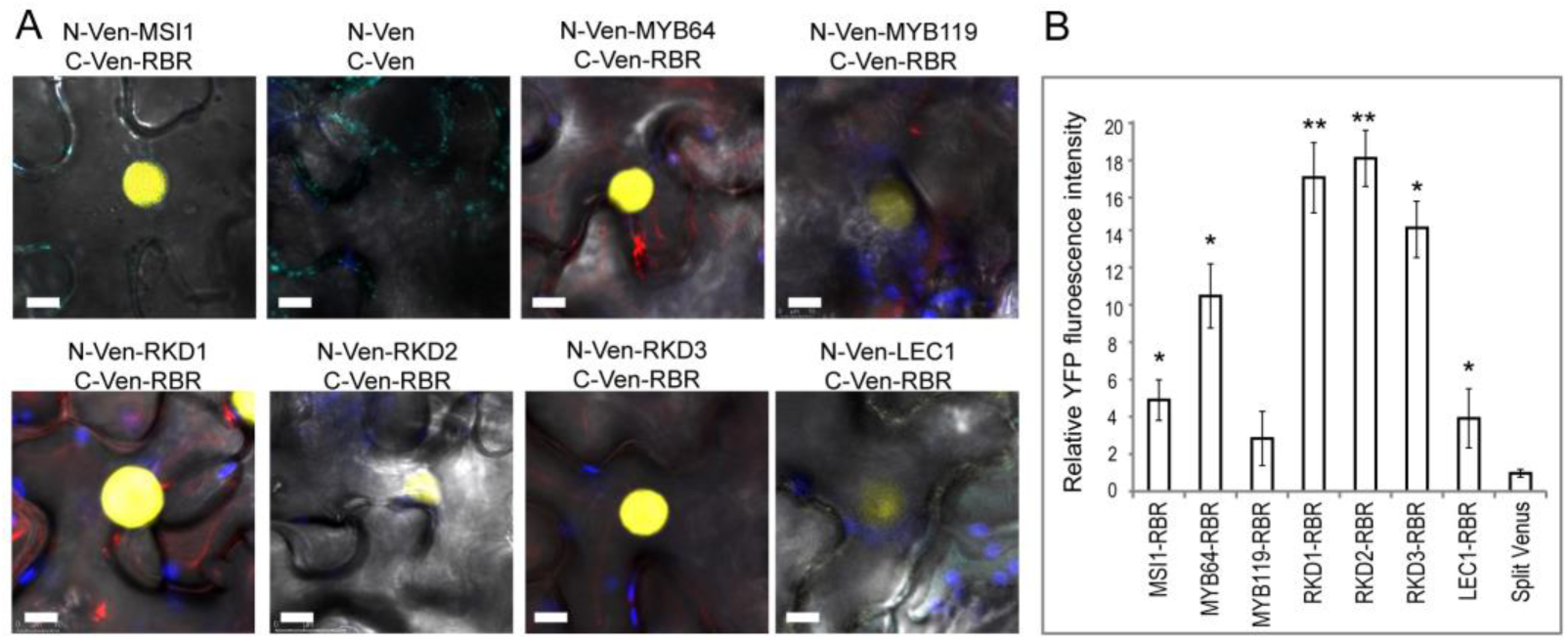
RBR physically interacts with MYB and RKD families of transcription factors. (**A**) Transient BiFC assay. RBR-MSI1 pair was used as a positive control [28], self-assembly of split-Venus as a negative control. (**B**) Quantification of BiFC interaction strength. Relative YFP fluorescence intensity in nuclei with background subtraction. C-Ven-RBR was tested with the respective N-Ven-protein fusions. Empty split Venus pair was used as a control in order to monitor unspecific background signals.

Whereas the above work established a subset of RBR interactors expressed in the egg cell, we wanted to supplement this work by building up a putative interaction map of the RBR egg cell network. We combined the available protein interaction datasets (Table S7), including those we identified in this work (Fig. 5, S4), and also we incorporated a subset of putative RBR interactors identified via a large-scale yeast-two-hybrid study that used a seedlings-specific *Arabidopsis* cDNA library as a bait (Gruissem lab, unpublished work in collaboration with Hybrigenics SA, Paris, France). We assumed each putative protein interactor to be present in the egg cell if the corresponding transcripts were previously identified to be a part of the *Arabidopsis* egg cell transcriptome [9]. The predicted network as depicted in Fig. S6 pinpointed that the putative RBR-centric egg cell interactome comprised of not only core cell cycle factors but also several differentiation and abiotic-stress associated nuclear factors.

## Discussion

### RBR & MYBs: cell cycle *versus* cell-cycle-independent mode of action?

Protein Retinoblastoma (pRB) and many MYB transcription factors are known cell-cycle regulators and onco-proteins in animal systems, and are part of evolutionarily ancient protein complexes [36–41]. Whereas pRB/RBR exists as a single or low copy number genes encoding conserved pocket proteins in animals and plants, the *MYB* genes that encode typical MYB domain proteins occur as single to multiple copies in animals. However, the plant MYB proteins comprise three subfamilies with some hundreds of proteins [42]. The MYBs reported here belong to a R2R3-type subfamily, which contains an additional homeo-domain. Protein interaction between RBR and MYB64/119 identified here suggests that RBR might form complexes with the plant MYBs. RBR-MYB protein interactions were previously reported for the cell cycle-related MYB3R-type proteins in *Arabidopsis* leaves [40] and for metazoan b-MYBs [43, 44]. Deregulation of both *RBR* and *MYB64/119* in *Arabidopsis* leads to super-numerary cells in the female gametophytes that are partially defective in establishing respective cell identities, which is reminiscent of cancer-like cell proliferation and oxidative cellular stress response. Female germline-specific requirement of RBR and MYBs in *Arabidopsis* and corresponding roles of their paralogues in mice oocytes [38, 45] illustrate how these dual modules might have retained their common reproductive function in evolution.

Three common aspects of the cell cycle regulation involving both RBR and MYB64 in *Arabidopsis* can be revisited. Firstly, the canonical cell cycle role of RB/RBR is to repress the transcription of E2F-regulated S-phase genes. Our ChIP data suggest that RBR might directly repress *MYB64* via an E2F canonical binding site in its promoter. Though *MYB64* has not been reported as a core cell cycle gene, *MYB64* transcripts were upregulated during the S-phase in an *Arabidopsis* suspension cell culture synchronized for cell cycle progression (Fig. S5) [46]. Therefore, we propose that *MYB64* might indeed be cell cycle regulated in the S-phase in *Arabidopsis*. The mechanism of MYB transcriptional regulation by RB appears to be evolutionarily ancient, as exemplified by similar cell-cycle regulation of the human Mybs [47, 48]. Secondly, depletion of RBR or MYB64/119 has strikingly similar effects and causes both rather cell-cycle-independent mis-establishment of cell identities and cell-cycle-dependent proliferation in the embryo sac, particularly of the egg cell [13, 22]. Thus, it is possible that both RBR and MYBs are essential for maintenance of the G2-phase pre-fertilization arrest and prevention of autonomous mitotic divisions of the egg cells. However, the mild upregulation of *MYB64* in *rbr-3* was unable to rescue egg cell function, indicating that complex interplay of both these proteins is necessary during female gamete development. A third aspect regarding an additional cell cycle role RBR and MYB64 concerns the mitotic phase. In the mature egg cells, RBR protein is quite low and MYB64 is rather abundant (Fig, 1C,I). Upon fertilization, zygotic expression of RBR declines even further, while the MYB64 signal is maintained at a similar level (Fig. 1C-E,I-K). The low abundance of RBR perhaps reflects an additional requirement of RBR in regulating mitotic division, similar to the M-phase-specific role of its paralog rblA in *Dictyostelium* [49]. Considering that MYB64 is detectable in the pre-mitotic zygote and elevation of its transcription around mitosis in the synchronized cell culture (Fig. S5), MYB64 might also function during the M-phase. Admittedly, we do not have an appropriate experimental setup with live plants yet to test for a) if and how RBR controls *MYB64* during the M-phase in early zygotic development; and b) if two-repeat R2R3-type MYB64 plays a role in M-phase in the egg cell similar to what the three-repeat 3R-type MYBs do in other plant tissues [40]. In addition, a cell-cycle-independent function of both RBR and MYBs [13, 22] from egg-to-zygote development will have to be investigated further.

It is also interesting to note that PRC2-specific repressive mark at the *MYB64* locus occurs in its gene body in the reproductive tissues, in contrast to the RBR-mediated repression of its promoter, indicating a rather complex regulation. Differential RBR binding between the two tested E2F binding sites in the *MYB64* promoter indicates its distinctive transcriptional regulation by RBR in reproductive versus sporophytic development. We also found RBR binding to *LEC1* and *WOX2* promoters, perhaps along with the H3K27me3 mark, illustrating possible combined RBR-PRC2 transcriptional regulation at their promoters during reproduction. Whereas *WOX2* seems to be directly repressed by RBR, and possibly also by the MYB64/119 and PRC2-mediated repression, RBR-MYBs function is necessary for maintaining *WOX8* expression. Together, RBR and MYB64/MYB119 play an important role in *WOX2/WOX8* balance in egg cell development, and probably also in ensuing zygote polarity establishment during the egg-to-zygotic reprogramming [8]. Therefore, RBR acts on promoters of a suite of transcription factors in the egg cell, while PRC2-dependent repression might play here an important parallel role. Cell-type specific data for the latter will have to be investigated in the future.

### RBR network mediates egg cell development and stress responses thereof

Previously, we have shown that the promoter activity of *RKD1* depends on intact RBR function in the egg cell [30]. Expression of *RKD*s and stress-related gene *LEC1* in the egg cell, a similar change of gene expression upon deregulation of RBR and MYB64/119 and upon stress, and their interaction with RBR, all indicate the underlying significance of this regulatory hub in plant reproduction. We propose that, unlike the RBR-MYB nodes described above, the network of RBR, RKDs and LEC1 is likely cell cycle-independent, but associated with maintenance of cellular homeostasis and stress response. The cross-regulation within the RKD clade is intriguing. Whether it is RBR or MYB-mediated cellular stress and cell differentiation, or most abiotic stress types tested here, *RKD3* was activated but *RKD2* was downregulated. *RKD3* repression in the wild-type is likely connected to H3K27me3 loading, and it is possible that RBR and PRC2 co-regulate this locus in a stress-responsive manner, and this dual transcriptional control is similar to regulation of other genes during seed maturation and early seedling development [31].

Environmental stress is a major denominator of evolution of sex and germline throughout the Eukaryotes [50], and the land plants in particular evolved across gradients of limiting water and increasing light conditions [51]. Surprisingly, we found that RBR represses a subset of egg cell-expressed stress-related genes, supporting its role in reproductive stress amelioration. Therefore, the stress-associated RBR-MYB-RKD-LEC1-(PRC2) transcription factor network that we uncovered here features a prominent higher-order regulatory mechanism that may underlie egg cell development and homeostasis in plants.

### Egg cell RBR network prevents parthenogenesis

The twin egg cell-like development observed in both the *rbr* and *myb* double mutants suggest that both RBR and MYB64/MYB119 are likely factors involved in preventing cell proliferation in the egg cell domain and that their deregulation could serve as a prerequisite for parthenogenesis. It is interesting to note that downregulation of MSI1, a member of RBR and PRC2 complexes, triggers early events of parthenogenesis [12]. Dominant-negative approach shows that deregulation of *RKDs* leads to formation of twin eggs and rare parthenogenesis-like events, supported by recent findings of similar events observed in knock-outs and knock-ins of the corresponding evolutionary homolog in *Marchantia*, and down-regulation of a *RKD2-* like gene in unreduced egg cells of *Boechera* at the onset of parthenogenesis [51–53]. Whereas the role of LEC1 during sexual egg cell development is not known, it is a crucial embryonic factor, overexpression of which is sufficient to induce somatic embryogenesis [18]. Interestingly, along with increase of embyogenic *LEC1* expression, abiotic stress induces abrogation of *RKD2*, *RKD3* derepression, and supernumerary egg production, supporting the general view that parthenogenesis evolved under stress conditions [54, 55].

Combining genetic, transcription and protein interaction data, we propose a model for RBR-centric transcription factor network in the egg cell (Fig. 6), which integrates stress amelioration and cellular homeostasis as inherent aspects of successful egg cell development coordinated by a subset of transcription factors. The proposed RBR-centric regulatory model and the putative hierarchical RBR-centric protein interaction network for the egg cell (Fig. S6) might help to dissect further intricate regulatory mechanisms involving stress and development.

**Fig. 6.**
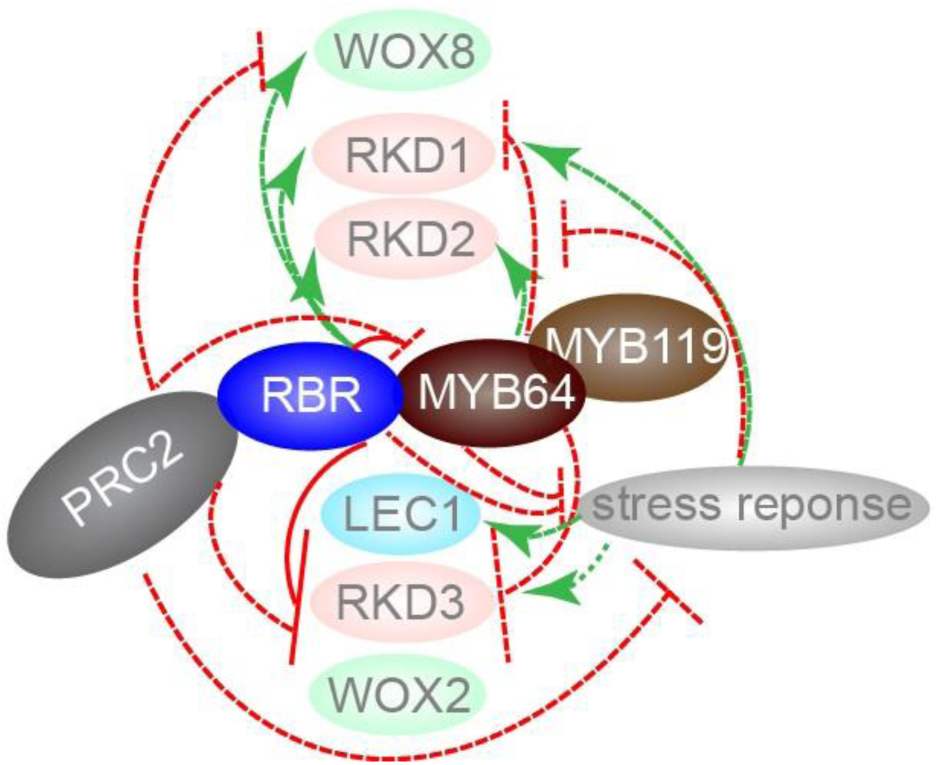
A model of cross-regulation between a subset of RBR-regulated transcription factors in the *Arabidopsis* egg cells. Illustrated is a schematic view of how interacting proteins RBR and MYBs commonly regulate a subset of transcription factor-encoding genes expressed in the egg cell. RBR/MYBs repress transcription of egg cell-specific *RKD3* and *WOX2*, and activate *RKD2 and WOX8*. RBR/MYB repress egg cell-expressed stress response genes, in particular *LEC1*, indicating their role in cellular homeostasis. RBR may mediate gene repression in concert with PRC2-specific repressive H3K27me3 mark.

### Materials and Methods

#### Plant material

Transgenic lines *rbr-3* [6, 24], *myb64-4*, *myb119-1*, *pMYB64::MYB64-GFP* [13], *pLEC1::GUS* [35] were described previously

#### Plasmid constructions

For stable *in planta* transformations, we used following binary vectors containing L1-L2 Gateway^®^ cassette, pK7WGF2 (VIB, Ghent) and p6N-GW (modified from the parent vector, DNA-Cloning-Service e.K., Hamburg). *RBR* coding and gene/genomic sequences and *RKD2* genic sequences were PCR-amplified directly from *Arabidopsis* accession Col-0 cDNA/DNA and were prepared as Gateway entry clones, as per manufacturer’s instructions (Thermo Fischer). *RKD2^DN^* sequence was cloned as a hybrid *RKD2* gene fused to *EAR* sequences of the *SUPERMAN* locus, generating a *RKD2^DN^* entry clone. A 2.2 Kbp *RBR* promoter (or) 1.3 Kbp *RKD2* promoter, 550 bp *pECA1.1* PCR-amplicon were cloned into the binary vectors by DNA ligation using T4-DNA ligase (Thermo Fischer). For transient *in vivo* protein-protein interactions we used binary vector pGWB601 (Nakagawa vectors, Addgene). For BiFC assembly, we used the *pUBI10*-driven Venus module sequences with very low self-assembly background signal described previously [56]. Portal clones of Split-Venus partner pairs were stacked by our recently developed cloning system “Byepass”, which utilized bacterial and yeast based endogenous recombination; cloning and vector details are presented elsewhere [57]. Terminal/binary clones were transformed into *Agrobacterium* via freeze-and-thaw method. Probes for *in situ* hybridization were PCR amplified as unique partial coding sequences of *MYB64*, *MYB119* and *EC1.1* from a cDNA pools from ovules, and cloned into an in-house expression vector.

#### Plant selection, cultivation and transformation

Surface-sterilized wild-type and transgenic seeds were germinated *in vitro* on MS half-strength plates without or with appropriate selection, subsequently transplanted into pots containing soil substrate, and cultivated in a long-day walk-in growth chamber conditions. Stable transformations of final agro-constructs were delivered into plants via floral-dip transformation [58]. A minimum of five independent transgenic lines were randomly chosen for genetic analysis of marker selection and seed set phenotyping; two representative lines were chosen for further analysis. Transient transformation of tobacco leaf mesophyll cells was achieved by *Agrobacterium*-mediated infiltration, as explained previously [59].

#### Abiotic stress induction

Stress treatments were given to soil-grown *Arabidopsis* plants. Flowering plants were exposed to stress conditions for one week before collection of pistils/ovules for down-stream analyses. For salt treatment, plants/pots were watered with 100mM NaCl every two days at 22°C; for elevated temperature treatment plants were placed at 27°C with sufficient watering; for drought treatment plants were minimally watered upon first signs of wilting at 22°C; control conditions were 22°C with normal watering regime every two days.

#### BiFC

Bimolecular Fluorescent Complementation assay was performed in young tobacco leaves upon transient agrobacterium-mediated transformation of BiFC constructs. RBR was fused at its N-terminus to C-Venus, and the tested interactors with N-Venus [56]. As negative controls, we used empty BiFC vector pair as well as C-Ven-RBR with empty N-Ven. Both combinations did not show meaningful fluorescence, as expected in BiFC experiments that used the improved parent vectors [56].

#### Microscopy

Fixed samples cleared in chloral hydrate for clearing analyses and/or those histochemically-stained for GUS detection [6] were observed under a Leica DMI6000 inverted microscope (Leica Microsystems) fitted with an Orca 4 camera (Hamamatsu). GUS staining was performed as in [22]. Confocal microscopy of Feulgen-stained samples [22] and/or live fluorescent samples were analysed under Zeiss LSM 780 (Carl Zeiss) or Leica SP8 (Leica Microsystems) confocal scanning laser microscopy platforms.

**mRNA *in situ* hybridization** was performed as described earlier [22]. Probes were prepared by *in vitro* transcription of appropriate template plasmids, and were hybridized on to 8 µm semi-thin sections of emasculated pistils containing mature ovules.

#### RNA extraction and cDNA synthesis

Minute ovule samples were pre-fixed in ethanol as described earlier [60] for all RT-PCR except for RNA-seq analysis ovules were scrapped out and snap-frozen immediately. It is important to note that the *rbr-3* mutant is homozygous lethal; therefore, plants heterozygous for *rbr-3* ubear only 50% ovules with haploid *rbr-3* embryo sacs that are encased by diploid integuments heterozygous for the same mutation [6,22,24]. *rbr-3* ovules were hand-picked as described in [28]. *myb64;myb119* ovules were pooled from homo-heterozygous double mutant plants [13]. Frozen tissues were ground in a tissue lyser (QiAGen), and the total RNA was prepared using RNA-Aqueous Micro kit and/or Trizol (Thermo Fischer), as per manufacturer’s instructions. Reverse transcription was performed on DNase I-treated samples using SuperScript IV First-Strand Synthesis System (Thermo Fischer).

#### mRNA-seq libraries and sequencing

Total RNA extracted from ovules of WT and mutant was quality-checked in a Bioanalyzer (Thermo Fischer). Purification of transcripts, library preparation and NGS sequencing were performed in a sequencing facility according to the routine pipeline (Fasteris, Switzerland). Paired-end sequencing generated approximately 100 bp per read in an Illumina HiSeq2000 platform.

#### Expression analysis based on mRNA-seq

The quality of raw reads was assessed using FastQC [61]. RNAseq samples from wild-type and *rbr-3/+* mature ovules were aligned to the *Arabidopsis* TAIR10 reference genome using Bowtie2 [62] with settings for sensitive mode. The number of uniquely mapped reads to each gene described in the reference genome annotation release of Araport11 [63] were counted using HTSeq [64]. Transcripts that were significantly differently expressed between wild-type and *rbr-3/+* ovules were identified using the NOISeq pipeline at a threshold of q > 0.95 [65]. The egg cell specific transcriptome between the wild-type and *rbr* mutant was derived from the overlap between the expressed transcripts in the wild-type and *rbr-3/+* ovules and the transcripts reported in a previous microarray from the egg cells [9]. Gene ontology (GO) enrichment analysis was done using BINGO [66] and visualized with Cytoscape [67].

**LexA-based yeast-two-hybrid growth assay** with RBR CDS was performed according to standard protocols. In brief, full-length coding sequences were cloned into modified pGilda bait (RBR) with 202-residue LexA domain, and pB24AD prey vectors (MATCHMAKER LexA Two-Hybrid System, Clontech), and transformed into high sensitivity yeast strain EGY48. Interactions were tested on synthetic complete (SC) yeast medium agar plates barring UHTL (Uracil, Histidine, Threonine, Leucine).

#### Protein interaction network analysis

Protein interaction network with egg cell enriched transcripts, transcription factor and down-stream stress related transcripts was created and visualized using GeneMANIA [68] and Cytoscape [67]. The primary networks were manually processed to remove non-significant and low confidence interactions to keep the network that had only physical and predicted interactions. Single nodes that did not have a direct link to RBR were removed, and the network was cropped to include the first node that linked RBR with a known stress associated protein. Additional links that created subnetworks from stress associated proteins included in the core network were also removed as these subnetworks did not add new information to link RBR to stress responses.

RBR and other interacting transcription factors are organized at the center followed by stress responsive genes at the outermost circle. The novel physical connections between RBR, MYB64, LEC1 and RKDs, discovered in the current study are shown in solid dark brown lines. The solid gray lines indicate published physical protein-protein interactions and the gray dashed-lines indicate published predicted protein-protein interactions. The intensity of the lines represents the significance of the interactions such as multiple independent studies. The white nodes represent proteins found to interact in published networks, but not enriched as transcripts in the deduced *rbr-3* egg cell transcriptome.

#### H3K27 trimethylation target identification

H3K27 trimethylation targets for the *Arabidopsis* flower tissues were obtained from the plantDHS database [69]. Upstream regions of the genes of interest in *Arabidopsis* genome [63] were searched at different window lengths to identify H3K27me3 targets. Finally, methylation targets in 500 upstream windows of the transcription start site (TSS) were reported as H3K27 trimethylation targets concentrated within this region.

#### Chromatin immuno-precipitation (ChIP)

RBR ChIP was performed on plants carrying *pRBR::GFP-RBR* in *rbr-3* background, *pEC::tagRFP-RBR*, and H3K27me3 ChIP on wild-type Col-0 plants. 3-week-old wild-type and *GFP-gRBR* seedlings, wild-type and *pRBR::GFP-RBR* inflorescences containing buds and open flowers before fertilization and gynoecia of wild-type and *pEC::tagRFP-RBR* unfertilized open flowers containing mature egg cells were collected. Egg-cell-targeted RBR ChIP was performed only for validation of a few fragments, as collection of material is extremely tedious and gynoecia contain a small proportion of egg cells resulting in a very low amount of bound DNA. Due to technical limitations in large-scale isolation of single egg cells required for H3K27me3 ChIP experiments, it was not possible to disentangle egg cell histone methylation patterns from the surrounding reproductive sporophytic tissues; therefore, we used mature unfertilized gynoecia. ChIP experiments were performed accordingly to the X-ChIP protocol as described in [59] using anti-GFP (Abcam, ab290), anti-RFP (AbCam, ab62341), anti-H3K27me3 (Millipore, #07-449) and anti-IgG (Abcam, ab6703) antibody.

#### Real-time qPCR

Both for RT-qPCR or ChIP-qPCR applications, SYBR Green assays were performed in QuantStudio5 Real-Time-PCR System (Thermo Fischer). A minimum of three biological replicates and two technical replicates were used in the experiments. The RT-qPCR data were normalized for expression of UBX domain-containing protein, *AT4G10790* [70]. For ChiP-qPCR, the values were normalized by the input, and background subtraction was performed for anti-GFP ChIP. Quantification of relative gene expression or DNA enrichment, and analyses of statistical inference using Student-t test were performed in Microsoft Excel 2010.

#### Statistical analysis

Fisher’s exact test http://graphpad.com/quickcalcs/contingency2/

## ACKNOWLEDGMENTS

## Supporting information

Supplemental Table 1,2,3,4, and all supplemental figure

## Acknowledgements

Acknowledged are funding from German Research Foundation (Emmy-Noether grant JO1001/1-1, DFG) and the Baden-Wuerttemberg State (LGFG) to A.J.J., the Max-Planck Society to F.T., National Science Foundation (NSF) to M.D, and Swiss National Science Foundation to W.G. We thank Gary Drews (University of Utah, USA) for kindly sharing transgenic material and advice, Claudia Casalongué (National University of Mar del Plata, Argentina) and Johan Peränen (University of Helsinki, Finland) for sharing yeast vectors, Marlene Zimmer (Heidelberg University) and Petra Tänzler (MPI Cologne) for technical help, Nora Mueller, Anika Rütz and Muhammad Tayyab (Heidelberg University) for help with cloning and plant work, Juan Mateo (University of Oviedo, Spain) for helpful suggestions regarding RNA-seq. Ivana Mesic and Claudia Sas (Thermo Scientific) for advice on real-time qPCR. We thank Nottingham Arabidopsis Stock Centre (NASC) for distributing transgenic seeds. Thanks are due to Sureshkumar Balasubramanian (Monash University, Australia) for critical comments on the manuscript.

## Author Contributions

A.J.J. designed and supervised the research. O.K., Pa.P., G.G., R.L., V.N, D.S.L., A.M.R., P.v.B. performed research. F.T. provided additional reagents and supervised specific experiments; O.K., Pr.P., J.S., C.W., Y.Z., W.G., F.T., M.D. and A.J.J. analysed data; M.D. supervised bioinformatic analysis; O.K. and A.J.J. wrote the paper with inputs from co-authors.

## Supporting Information

Table S1. A hemizygous transgene *GFP-gRBR* fully restores fertility of *rbr-3* gametophytes in *rbr-3/RBR; GFP-gRBR^h^*.

Table S2. Progeny test confirms ‘two independent loci’ complementation of rbr-3 allele with GFP-gRBR in rbr-3/RBR; GFP-gRBR^h^ background. Note that offspring was scored based on antibiotic resistance of rbr-3 T-DNA.

Table S3. A hemizygous transgene *pEC-gRBR* partially restores fertility of *rbr-3* gametophytes in *rbr-3/RBR; pEC1-gRBR^h^* background.

Table S4. Progeny test confirms partial ‘two independent loci’ complementation of rbr-3 allele with pEC-gRBR in rbr-3/RBR; pEC-gRBR^h^ background. Note that offspring was scored based on antibiotic resistance of rbr-3 T-DNA.

Table S5. Previously validated egg cell expressed transcripts showing deregulation in *rbr-3* ovule transcriptome

Table S6. List of RBR-regulated transcription factors, and stress and stimulus-responsive transcripts in the egg cell, subtracted from the ovule transcriptomes

Table S7. List of egg cell-expressed RBR interactors used for building protein interaction network.

**Fig. S1. Overall gene ontology enrichment of stress and stimulus responsive genes and transcription factors in *rbr-3* egg cells.** Gene ontology clustering for abiotic stress and stimulus enriched (*A*) or depleted (*B*) in *rbr-3* ovules, and enriched (*C*) or depleted (*D*) in *rbr-3* egg cells.

**Fig. S2. *MYB64* and *MYB119* transcripts in the mature embryo sac.** mRNA *in situ* hybridization: (A) Sense (*S*) probes shows no signal, while anti-sense probes (*AS*) detect (B) *MYB64* and (C) *MYB119* mRNA in the mature embryo sac, and specifically in the egg cell. Scale bar=20µm.

**Fig. S3. Chromatin immunoprecipitation in reproductive tissues in comparison to vegetative stages in *Arabidopsis.*** (A) ChIP for RBR or (B) for the PRC2-specific H3K27me3 binding. Relative real-time qPCR data normalized by input. *PCNA* locus was used as a positive control for RBR binding but negative for H3K27me3. Significant difference is indicated between seedling and inflorescence tissues: **α ≤ 0.01; *α ≤ 0.05.

**Fig. S4. RBR interacts with egg-cell expressed transcription factors.** (A) RBR protein-protein interactions identified by LexA-based yeast-two-hybrid assay. (B) Relative nuclei fluorescence intensity in BiFC assay with background subtraction. C-Ven-RBR was tested with the respective N-Ven-protein fusions.

**Fig. S5. Cell-cycle-dependent expression of *MYB64*.** *MYB64* transcript signals change with cell cycle progression in synchronized *Arabidopsis* cell culture in an opposite manner to *RBR* [46].

**Fig. S6. RBR connects transcriptional regulation and stress response shared by PRC2.** A protein interaction network connecting RBR, cell cycle, transcription factors and stress-responsive genes. Transcription factors enriched in *rbr-3* egg cell transcriptome connect to stress responsive

