## Supplemental Table 5 for "A RETINOBLASTOMA-RELATED transcription factor network governs egg cell differentiation and stress response in *Arabidopsis*"

Supplementary Table 5. Previously validated egg cell expressed transcripts showing deregulation in *rbr3* ovule transcriptome

| Locus | Gene Description | EC expression | Pubmed ID | DETs | Validated |
|---|---|---|---|---|---|
| **Upregulated genes** | | | | | |
| At4g01970 | RAFFINOSE SYNTHASE 4, RS4 | ERF | 17915010 | up | |
| At5g56200 | C2H2 type zinc finger transcription factor family | PRF | 20550711 | up | |
| At5g59340 | WUSCHEL RELATED HOMEOBOX 2, WOX2 | ISH | 14711878 | up | yes |
| At5g62270 | GAMETE CELL DEFECTIVE 1, ribosomal protein L20 | PRF | 23085019 | up | |
| At3g62320 | unknown | PRF | 21457369 | up | |
| At1g66610 | TRAF-like superfamily protein | PRF | 21457369 | up | |
| AT1G08130 | DNA ligase | TLRF | 20023162 | up, NS | |
| AT5G49160 | DNA METHYLTRANSFERASE 1, MET1 | ISH | 23146178 | up, NS | yes |
| At1g80370 | CYCLIN A2;4 | ERF | 17915010 | up, NS | |
| At5g16850 | TELOMERASE REVERSE TRANSCRIPTASE | ISH | 20226671 | up, NS | |
| At4g02060 | MCM7, PRL, PROLIFERA | ERF | 7754372 | up, NS | |
| **Down-regulated genes** | | | | | |
| At1g78940 | Kinase | ISH | 17915010 | down | |
| At1g76750 | EC1.1 | ISH | 23180860 | down | yes |
| At5g45980 | WUSCHEL RELATED HOMEOBOX 8, WOX8 | ISH | 14711878 | down | yes |
| At5g14620 | DOMAINS REARRANGED METHYLASE 2, DRM2 | TLRF | 22940470 | down | |
| At4g20050 | pectin lyase, QUARTET 3 | PRF | 17559508 | down, NS | |
| At5g55820 | WYRD nner centromere protein (INCENP) | ISH | 21752930 | down, NS | |
| At2g41500 | LACHESIS, Splicing factor | ISH | 17326723 | down, NS | |
| At1g06220 | CLOTHO, GAMETOPHYTE FACTOR 1, GFA1, Splicing factor | ISH | 18702672 | down, NS | |
| AT1g60530 | DYNAMIN RELATED PROTEIN 4A, DRP4A | PRF | 21457369 | down, NS | |
| At5g55490 | GAMETE EXPRESSED 1, GEX1 | PRF | 21831199 | down, NS | |

ISH - mRNA in situ hybridization
PRF-promoter reporter fusion
ERF - Enhancer reporter Fusion
TLRF -Translational reporter fusion
NS - not statistically significant in our analysis
DETs- differentially expressed transcripts
