## Supplemental Table 7 for "A RETINOBLASTOMA-RELATED transcription factor network governs egg cell differentiation and stress response in *Arabidopsis*"

Supplementary Table S7. List of egg cell-expressed RBR interactors used for building protein interaction network.

| GeneIDs | Gene name | Gene description | Normalized read count for H3K27M3 target |
|---|---|---|---|
| AT1G05180 | AXR1 | NAD(P)-binding Rossmann-fold superfamily protein | 1 |
| AT1G08130 | LIG1 | DNA ligase 1 | 1 |
| AT1G08970 | NF-YC9 | nuclear factor Y, subunit C9 | 1 |
| AT1G13440 | GAPC2 | glyceraldehyde-3-phosphate dehydrogenase C2 | 2 |
| AT1G15570 | CYCA2;3 | CYCLIN A2;3 | 1 |
| AT1G20200 | EMB2719 | PAM domain (PCI/PINT associated module) protein | 1 |
| AT1G21970 | LEC1 | Histone superfamily protein | 5 |
| AT1G30580 | AT1G30580 | GTP binding protein | 2 |
| AT1G45000 | AT1G45000 | AAA-type ATPase family protein | 1 |
| AT1G47870 | ATE2F2 | winged-helix DNA-binding transcription factor family protein | 3 |
| AT1G49240 | ACT8 | actin 8 | 0 |
| AT1G54100 | ALDH7B4 | aldehyde dehydrogenase 7B4 | 1 |
| AT1G54830 | NF-YC3 | nuclear factor Y, subunit C3 | 1 |
| AT1G56450 | PBG1 | 20S proteasome beta subunit G1 | 6 |
| AT1G60490 | VPS34 | vacuolar protein sorting 34 | 1 |
| AT1G64790 | ILA | ILITYHIA | 4 |
| AT1G66340 | ETR1 | Signal transduction histidine kinase, hybrid-type, ethylene sensor | 2 |
| AT1G66750 | CAK4 | CDK-activating kinase 4 | 1 |
| AT2G04520 | AT2G04520 | Nucleic acid-binding, OB-fold-like protein | 2 |
| AT2G28190 | CSD2 | copper/zinc superoxide dismutase 2 | 6 |
| AT2G36010 | E2F3 | E2F transcription factor 3 | 3 |
| AT2G37060 | NF-YB8 | nuclear factor Y, subunit B8 | 3 |
| AT2G44530 | AT2G44530 | Phosphoribosyltransferase family protein | 1 |
| AT2G45640 | SAP18 | SIN3 associated polypeptide P18 | 3 |
| AT3G09480 | AT3G09480 | Histone superfamily protein | 1 |
| AT3G11250 | AT3G11250 | Ribosomal protein L10 family protein | 1 |
| AT3G11630 | AT3G11630 | Thioredoxin superfamily protein | 3 |
| AT3G12280 | RBR1 | retinoblastoma-related 1 | 1 |
| AT3G17210 | HS1 | heat stable protein 1 | 1 |
| AT3G18780 | ACT2 | actin 2 | 11 |
| AT3G22845 | AT3G22845 | emp24/gp25L/p24 family/GOLD family protein | 1 |
| AT3G46970 | PHS2 | alpha-glucan phosphorylase 2 | 1 |
| AT3G48750 | CDC2 | cell division control 2 | 2 |
| AT3G53340 | NF-YB10 | nuclear factor Y, subunit B10 | 1 |
| AT3G55280 | RPL23AB | ribosomal protein L23AB | 1 |
| AT3G58180 | AT3G58180 | ARM repeat superfamily protein | 3 |
| AT3G59380 | FTA | farnesyltransferase A | 1 |
| AT4G01370 | MPK4 | MAP kinase 4 | 7 |
| AT4G11600 | GPX6 | glutathione peroxidase 6 | 2 |
| AT4G24820 | AT4G24820 | 26S proteasome regulatory subunit Rpn7 | 2 |
| AT4G37630 | CYCD5;1 | cyclin d5;1 | 5 |
| AT5G02450 | AT5G02450 | Ribosomal protein L36e family protein | 5 |
| AT5G02470 | DPA | Transcription factor DP | 2 |
| AT5G03415 | DPB | Transcription factor DP | 1 |

| | | | |
|---|---|---|---|
| AT5G06290 | 2-Cys Prx B | 2-cysteine peroxiredoxin B | 7 |
| AT5G09500 | AT5G09500 | Ribosomal protein S19 family protein | 4 |
| AT5G16970 | AER | alkenal reductase | 4 |
| AT5G17310 | UGP2 | UDP-glucose pyrophosphorylase 2 | 1 |
| AT5G18620 | CHR17 | chromatin remodeling factor17 | 1 |
| AT5G22220 | E2F1 | E2F transcription factor 1 | 2 |
| AT5G27620 | CYCH;1 | cyclin H;1 | 6 |
| AT5G40280 | ERA1 | Prenyltransferase family protein | 18 |
| AT5G40580 | PBB2 | 20S proteasome beta subunit PBB2 | 1 |
| AT5G54640 | RAT5 | Histone superfamily protein | 2 |
| AT5G55070 | AT5G55070 | Dihydrolipoamide succinyltransferase | 2 |
| AT5G11050 | MYB64 | myb domain protein 64 | 1 |
| AT1G18790 | RKD1 | RWP-RK domain-containing protein | 18 |
| AT1G74480 | RKD2 | RWP-RK domain-containing protein | 3 |
| AT5G66990 | RKD3 | RWP-RK domain-containing protein | 33 |

#H3K27M3 target was determined based on normalized read count value present in PlantDHS database for respective gene IDs.
