## Supplemental Table 6.1 for "A RETINOBLASTOMA-RELATED transcription factor network governs egg cell differentiation and stress response in *Arabidopsis*"

**SI Appendix Table S6.1** List of RBR-regulated transcription factors, and stress and stimulus-responsive transcripts in the egg cell, subtracted from the ovule transcriptomes (A) Transcription Factors

| *Arabidopsis* Locus ID | Gene Model Description | TF Family Name |
|---|---|---|
| **I. Enriched in *rbr-3*** | | |
| AT1G08970 | nuclear factor Y%2C subunit C9 | CCAAT-HAP5 |
| AT1G14410 | ssDNA-binding transcriptional regulator | Whirly |
| AT1G14510 | alfin-like 7 | Alfin |
| AT1G22070 | transcription factor TGA3 | bZIP |
| AT1G32700 | PLATZ transcription factor family protein | PLATZ |
| AT1G34370 | C2H2 and C2HC zinc fingers superfamily protein | C2H2 |
| AT1G42990 | basic region/leucine zipper motif 60 | bZIP |
| AT1G63490 | transcription factor jumonji (jmjC) domain-containing protein | JUMONJI |
| AT2G02160 | CCCH-type zinc finger family protein | C3H |
| AT2G17410 | ARID/BRIGHT DNA-binding domain-containing protein | ARID |
| AT2G24430 | NAC domain containing protein 38 | NAC |
| AT2G27050 | ETHYLENE-INSENSITIVE3-like 1 | EIL |
| AT2G34720 | nuclear factor Y%2C subunit A4 | CCAAT-HAP2 |
| AT2G35430 | Zinc finger C-x8-C-x5-C-x3-H type family protein | C3H |
| AT2G36010 | E2F transcription factor 3 | E2F-DP |
| AT2G39830 | DA1-related protein 2 | LIM |
| AT3G07670 | Rubisco methyltransferase family protein | PcG |
| AT3G10500 | NAC domain containing protein 53 | NAC |
| AT3G10760 | Homeodomain-like superfamily protein | GARP-G2-like |
| AT3G12130 | KH domain-containing protein / zinc finger (CCCH type) family protein | C3H |
| AT3G59580 | Plant regulator RWP-RK family protein | Nin-like |
| AT4G17750 | heat shock factor 1 | HSF |
| AT4G24470 | GATA-type zinc finger protein with TIFY domain-containing protein | ZIM |
| AT4G30935 | WRKY DNA-binding protein 32 | WRKY |
| AT4G31610 | Transcriptional factor B3 family protein | ABI3-VP1 |
| AT5G02810 | pseudo-response regulator 7 | C2C2-CO-like |
| AT5G06770 | KH domain-containing protein / zinc finger (CCCH type) family protein | C3H |
| AT5G08520 | Duplicated homeodomain-like superfamily protein | MYB |
| AT5G11050 | myb domain protein 64 | MYB |
| AT5G27610 | protein ALWAYS EARLY 1 | MYB-related |
| AT5G51910 | TCP family transcription factor | TCP |
| AT5G59820 | C2H2-type zinc finger family protein | C2H2 |
| AT1G21000 | PLATZ transcription factor family protein | PLATZ |
| AT1G25470 | AP2 domain-containing transcription factor family protein | AP2-EREBP |
| AT1G47270 | tubby like protein 6 | TLP |
| AT1G49950 | telomere repeat binding factor 1 | MYB-related |
| AT1G68920 | basic helix-loop-helix (bHLH) DNA-binding superfamily protein | bHLH |
| AT1G74950 | TIFY domain/Divergent CCT motif family protein | ZIM |
| AT2G24500 | Zinc finger protein 622 | C2H2 |
| AT2G28510 | Dof-type zinc finger DNA-binding family protein | C2C2-Dof |
| AT2G32700 | LEUNIG-like protein | LUG |
| AT2G47680 | zinc finger (CCCH type) helicase family protein | C3H |

| | | |
|---|---|---|
| AT3G06410 | Zinc finger C-x8-C-x5-C-x3-H type family protein | C3H |
| AT3G10800 | Basic-leucine zipper (bZIP) transcription factor family protein | bZIP |
| AT3G58680 | multiprotein bridging factor 1B | MBF1 |
| AT4G17710 | homeodomain GLABROUS 4 | HB |
| AT5G64060 | NAC domain containing protein 103 | NAC |
| AT5G64220 | Calmodulin-binding transcription activator protein with CG-1 and Ankyrin domain | CAMTA |
| AT2G35550 | basic pentacysteine 7 | BBR-BPC |
| AT2G36480 | pre-mRNA cleavage complex 2 Pcf11-like protein | C2H2 |
| AT3G24010 | RING/FYVE/PHD zinc finger superfamily protein | PHD |
| AT3G25730 | ethylene response DNA binding factor 3 | AP2-EREBP |
| AT3G62240 | RING/U-box superfamily protein | C2H2 |
| AT5G05610 | alfin-like 1 | Alfin |
| AT5G46910 | Transcription factor jumonji (jmj) family protein / zinc finger (C5HC2 type) family protein | JUMONJI |
| AT5G52660 | Homeodomain-like superfamily protein | MYB-related |
| AT5G59340 | WUSCHEL related homeobox 2 | HB |
| AT2G40950 | Basic-leucine zipper (bZIP) transcription factor family protein | bZIP |
| AT5G47390 | myb-like transcription factor family protein | MYB-related |
| AT1G72050 | transcription factor IIIA | C2H2 |
| AT3G53680 | Acyl-CoA N-acyltransferase with RING/FYVE/PHD-type zinc finger domain-containing protein | PHD |
| AT1G34190 | NAC domain containing protein 17 | NAC |
| AT5G38840 | SMAD/FHA domain-containing protein | FHA |
| AT5G58080 | response regulator 18 | GARP-G2-like |
| AT1G77950 | AGAMOUS-like 67 | MADS |
| AT5G18680 | tubby like protein 11 | TLP |

**II. depleted in *rbr-3***

| | | |
|---|---|---|
| AT1G12860 | basic helix-loop-helix (bHLH) DNA-binding superfamily protein | bHLH |
| AT1G50620 | RING/FYVE/PHD zinc finger superfamily protein | PHD |
| AT1G51140 | basic helix-loop-helix (bHLH) DNA-binding superfamily protein | bHLH |
| AT1G58220 | Homeodomain-like superfamily protein | MYB-related |
| AT2G21530 | SMAD/FHA domain-containing protein | FHA |
| AT2G28810 | Dof-type zinc finger DNA-binding family protein | C2C2-Dof |
| AT2G32930 | zinc finger nuclease 2 | C3H |
| AT2G36930 | zinc finger (C2H2 type) family protein | C2H2 |
| AT2G37060 | nuclear factor Y%2C subunit B8 | CCAAT-HAP3 |
| AT2G47890 | B-box type zinc finger protein with CCT domain-containing protein | C2C2-CO-like |
| AT3G16500 | phytochrome-associated protein 1 | AUX-IAA |
| AT3G17860 | jasmonate-zim-domain protein 3 | ZIM |
| AT3G53340 | nuclear factor Y%2C subunit B10 | CCAAT-HAP3 |
| AT4G14560 | indole-3-acetic acid inducible | AUX-IAA |
| AT4G17230 | SCARECROW-like 13 | GRAS |
| AT4G25210 | DNA-binding storekeeper protein-related transcriptional regulator | GeBP |
| AT4G26640 | WRKY family transcription factor family protein | WRKY |
| AT4G27910 | SET domain protein 16 | PHD |
| AT4G36920 | Integrase-type DNA-binding superfamily protein | AP2-EREBP |
| AT5G09790 | TRITHORAX-RELATED PROTEIN 5 | PHD |
| AT5G12440 | CCCH-type zinc fingerfamily protein with RNA-binding domain-containing protein | C3H |

| | | |
|---|---|---|
| AT5G14960 | DP-E2F-like 2 | E2F-DP |
| AT5G18550 | Zinc finger C-x8-C-x5-C-x3-H type family protein | C3H |
| AT5G25475 | AP2/B3-like transcriptional factor family protein | ABI3-VP1 |
| AT5G42520 | basic pentacysteine 6 | BBR-BPC |
| AT5G45300 | beta-amylase 2 | BES1 |
| AT5G45980 | WUSCHEL related homeobox 8 | HB |
| AT5G60410 | DNA-binding protein with MIZ/SP-RING zinc finger%2C PHD-finger and SAP domain-containing protein | PHD |
| AT3G23050 | indole-3-acetic acid 7 | AUX-IAA |
| AT3G61150 | homeodomain GLABROUS 1 | HB |
| AT4G38000 | DNA binding with one finger 4.7 | C2C2-Dof |
| AT5G61190 | putative endonuclease or glycosyl hydrolase with C2H2-type zinc finger domain-containing protein | C2H2 |
| AT1G76510 | ARID/BRIGHT DNA-binding domain-containing protein | ARID |
| AT4G02560 | Homeodomain-like superfamily protein | HB |
| AT4G15180 | SET domain protein 2 | PcG |
| AT4G32880 | homeobox-leucine zipper protein ATHB-8 | HB |
| AT1G54060 | 6B-interacting protein 1-like 1 | Trihelix |
| AT4G31060 | Integrase-type DNA-binding superfamily protein | AP2-EREBP |
| AT1G15580 | indole-3-acetic acid inducible 5 | AUX-IAA |

#H3K27M3 target was determined based on normalized read count value present in PlantDHS database for respective gene IDs.

### RBR/E2F binding site was predicted running FIMO algorithm using published E2F binding motifs as templates

| H3K27M3 target | RBR/E2F binding site | stress responsive | | TF description |
|---|---|---|---|---|
| Yes/No | Yes/No | P-value | Yes/No | |
| Yes | Yes | 2.55E-05 | No | CCAAT-box binding transcription factor Hap5a, putative chr1:2882546-2884285 FORWARD |
| Yes | Yes | 9.65E-08 | Yes | DNA-binding protein-related, similar to DNA-binding protein p24 GI:9651810 from (Solanum |
| Yes | Yes | 7.48E-05 | No | PHD finger family protein, contains Pfam domain, PF00628: PHD-finger chr1:4961882-4964 |
| Yes | Yes | 5.02E-05 | Yes | bZIP family transcription factor (TGA3), identical to transcription factor GI:304113 from (Aral |
| Yes | Yes | 7.51E-05 | No | zinc-binding family protein, similar to zinc-binding protein (Pisum sativum) GI:16117799; cor |
| Yes | Yes | 2.99E-05 | No | similar to zinc finger (C2H2 type) family protein [Arabidopsis thaliana] (TAIR:At5g22890.1); s |
| Yes | No | #N/A | Yes | AtbZIP60 consists of a bZIP DNA binding domain followed by a putative transmembrane dom |
| Yes | Yes | 9.48E-05 | No | transcription factor jumonji (jmjC) domain-containing protein, similar to PLU-1 protein (GI:4 |
| Yes | No | #N/A | No | zinc finger (CCCH-type) family protein, contains Pfam domain, PF00642: Zinc finger C-x8-C-x |
| Yes | Yes | 9.2E-06 | No | ARID/BRIGHT DNA-binding domain-containing protein, contains Pfam profile PF01388: ARID |
| Yes | Yes | 6.07E-06 | No | no apical meristem (NAM) family protein, contains Pfam PF02365: No apical meristem (NAM |
| Yes | Yes | 5.18E-05 | Yes | ethylene-insensitive3-like1 (EIL1), identical to ethylene-insensitive3-like1 GI:2224927 from ( |
| Yes | Yes | 4.02E-05 | No | CCAAT-binding transcription factor (CBF-B/NF-YA) family protein, contains Pfam profile: PF0 |
| Yes | Yes | 1.58E-05 | No | zinc finger (CCCH-type) family protein, contains Pfam domain, PF00642: Zinc finger C-x8-C-x |
| Yes | Yes | 2.91E-05 | No | E2F transcription factor-3 (E2F3), identical to E2F transcription factor-3 E2F3 (Arabidopsis th |
| Yes | Yes | 1.38E-10 | No | LIM domain-containing protein, contains Pfam profile PF00412: LIM domain chr2:16626855 |
| Yes | Yes | 2.06E-05 | No | SET domain-containing protein, similar to ribulose-1,5-bisphosphate carboxylase/oxygenase |
| Yes | Yes | 0.000028 | No | no apical meristem (NAM) family protein, similar to to NAC2 (GI:645671) (Arabidopsis thaliar |
| Yes | Yes | 6.31E-06 | No | myb family transcription factor, contains Pfam profile: PF00249 myb-like DNA-binding doma |
| Yes | Yes | 4.06E-05 | No | KH domain-containing protein / zinc finger (CCCH type) family protein chr3:3864158-38665 |
| Yes | Yes | 5.64E-05 | No | similar to RWP-RK domain-containing protein [Arabidopsis thaliana] (TAIR:At2g43500.1); sim |
| Yes | Yes | 6.08E-07 | Yes | heat shock factor protein 1 (HSF1) / heat shock transcription factor 1 (HSTF1), identical to he |
| Yes | Yes | 7.4E-06 | No | zinc finger (GATA type) protein ZIM (ZIM), identical to zinc-finger protein expressed in Inflore |
| Yes | Yes | 2.45E-05 | No | WRKY family transcription factor, contains Pfam profile: PF03106 WRKY DNA -binding domai |
| Yes | Yes | 2.61E-06 | No | transcriptional factor B3 family protein / reproductive meristem protein 1 (REM1), similar to |
| Yes | Yes | 2.72E-06 | No | pseudo-response regulator 7 (APRR7), identical to pseudo-response regulator 7 GI:1028100 |
| Yes | Yes | 4.8E-06 | No | KH domain-containing protein / zinc finger (CCCH type) family protein, contains Pfam domai |
| Yes | Yes | 4.05E-05 | No | myb family transcription factor, contains Pfam profile: PF00249 myb-like DNA-binding doma |
| Yes | Yes | 6.99E-06 | No | myb family transcription factor, contains Pfam profile: PF00249 myb-like DNA binding doma |
| Yes | Yes | 3.47E-05 | No | similar to myb family transcription factor [Arabidopsis thaliana] (TAIR:At3g05380.1); similar |
| Yes | Yes | 1.42E-06 | No | TCP family transcription factor, putative, similar to PCF2 ((GI:2580440) Oryza sativa) chr5:21 |
| Yes | Yes | 3.58E-06 | Yes | zinc finger (C2H2 type) family protein (ZAT12), identical to zinc finger protein ZAT12 (Arabid |
| Yes | No | #N/A | No | zinc-binding family protein, similar to zinc-binding protein (Pisum sativum) GI:16117799; cor |
| Yes | Yes | 0.000026 | No | encodes a member of the ERF (ethylene response factor) subfamily B-6 of ERF/AP2 transcrip |
| Yes | Yes | 0.000033 | No | F-box family protein / tubby family protein, contains Pfam profiles: PF00646 F-box domain, F |
| Yes | Yes | 1.38E-05 | Yes | DNA-binding protein, putative, contains similarity to DNA-binding protein PcMYB1 (Petroseli |
| Yes | Yes | 0.000072 | No | similar to basic helix-loop-helix (bHLH) family protein [Arabidopsis thaliana] (TAIR:At1g2626 |
| Yes | Yes | 1.17E-05 | Yes | expressed protein chr1:28152236-28154055 REVERSE Aliases: F25A4.8, F25A4_8 |
| Yes | Yes | 1.56E-05 | No | zinc finger (C2H2 type) family protein, contains Pfam profile: PF00096 zinc finger, C2H2 type |
| Yes | Yes | 1.29E-05 | No | Dof-type zinc finger domain-containing protein, similar to elicitor-responsive Dof protein ERI |
| Yes | Yes | 6.99E-05 | No | similar to WD-40 repeat family protein (LEUNIG) [Arabidopsis thaliana] (TAIR:At4g32551.1); |
| Yes | Yes | 0.00006 | No | zinc finger (CCCH type) helicase family protein, similar to SP:Q28141 ATP-dependent RNA he |

| | | | | |
|---|---|---|---|---|
| Yes | Yes | 6.07E-06 | No | similar to zinc finger (CCCH-type) family protein [Arabidopsis thaliana] (TAIR:At5g18550.1); s |
| Yes | Yes | 4.87E-05 | Yes | bZIP transcription factor family protein, contains Pfam profile: PF00170 bZIP transcription fa |
| Yes | Yes | 4.14E-05 | No | ethylene-responsive transcriptional coactivator, putative, similar to ethylene-responsive tra |
| Yes | Yes | 7.39E-05 | No | homeobox-leucine zipper family protein / lipid-binding START domain-containing protein, si |
| Yes | Yes | 2.21E-05 | No | no apical meristem (NAM) family protein, similar to NAC1 (GI:7716952) {Medicago truncatul |
| Yes | Yes | 2.72E-06 | No | similar to ethylene-responsive calmodulin-binding protein, putative (SR1) [Arabidopsis thalia |
| Yes | Yes | 3.06E-05 | No | similar to expressed protein [Arabidopsis thaliana] (TAIR:At1g14685.3); similar to expressed |
| Yes | No | #N/A | No | zinc finger (C2H2-type) family protein, weak similarity to S-locus protein 4 (GI:6069478) (Bra |
| Yes | No | #N/A | No | PHD finger family protein, contains Pfam profile: PF00628 PHD-finger chr3:8675965-867822 |
| Yes | Yes | 1.61E-05 | No | AP2 domain-containing transcription factor, putative, contains Pfam profile: PF00847 AP2 do |
| Yes | Yes | 1.01E-05 | No | zinc finger (C2H2 type) family protein, contains Pfam PF00096: Zinc finger, C2H2 type chr3:2 |
| Yes | Yes | 0.000011 | No | PHD finger family protein, contains Pfam domain, PF00628: PHD-finger chr5:1676941-1679 |
| Yes | Yes | 8.05E-05 | No | transcription factor jumonji (jmj) family protein, contains Pfam domains PF02375: jmjN dom |
| Yes | Yes | 4.71E-06 | Yes | myb family transcription factor, contains PFAM profile: PF00249 myb-like DNA-binding doma |
| Yes | Yes | 2.69E-05 | No | similar to homeobox-leucine zipper transcription factor family protein [Arabidopsis thaliana] |
| Yes | Yes | 8.47E-05 | Yes | bZIP transcription factor family protein, similar to AtbZIP transcription factor GI:17065880 fr |
| Yes | Yes | 1.04E-05 | Yes | myb family transcription factor, contains Pfam profile: PF00249 myb-like DNA-binding doma |
| Yes | Yes | 2.43E-05 | No | zinc finger (C2H2 type) family protein, contains multiple zinc finger domains: PF00096: Zinc f |
| Yes | Yes | 2.14E-07 | No | PHD finger transcription factor, putative, predicted proteins, Arabidopsis thaliana chr3:1990 |
| Yes | Yes | 4.03E-05 | No | no apical meristem (NAM) family protein, contains Pfam PF02365: No apical meristem (NAM |
| Yes | Yes | 2.28E-05 | No | forkhead-associated domain-containing protein / FHA domain-containing protein, related to |
| Yes | Yes | 0.000038 | No | two-component responsive regulator family protein / response regulator family protein, con |
| Yes | Yes | 3.71E-05 | No | similar to MADS-box family protein [Arabidopsis thaliana] (TAIR:At1g22130.1); similar to put |
| Yes | Yes | 4.14E-05 | No | F-box family protein / tubby family protein, similar to phosphodiesterase (GI:467578) (Mus r |

| | | | | |
|---|---|---|---|---|
| Yes | Yes | 3.66E-05 | Yes | basic helix-loop-helix (bHLH) family protein / F-box family protein, contains Pfam profiles: PF |
| Yes | Yes | 1.18E-05 | No | PHD finger family protein, contains Pfam domain, PF00628: PHD-finger chr1:18752205-187 |
| Yes | Yes | 2.16E-05 | No | basic helix-loop-helix (bHLH) family protein, contains Pfam profile: PF00010 helix-loop-helix |
| Yes | Yes | 3.96E-05 | No | myb family transcription factor, contains Pfam profile: PF00249: Myb-like DNA-binding doma |
| Yes | No | #N/A | No | forkhead-associated domain-containing protein / FHA domain-containing protein chr2:9226 |
| Yes | No | #N/A | No | Dof-type zinc finger domain-containing protein, similar to zinc finger protein OBP2 GI:50593 |
| Yes | Yes | 7.29E-05 | No | zinc finger (CCCH-type) family protein, contains Pfam domain, PF00642: Zinc finger C-x8-C-x |
| Yes | Yes | 8.42E-05 | No | zinc finger (C2H2 type) family protein, contains Prosite PS00028: Zinc finger, C2H2 type, dom |
| Yes | Yes | 4.05E-05 | No | similar to CCAAT-box binding transcription factor, putative [Arabidopsis thaliana] (TAIR:At3g |
| Yes | Yes | 1.69E-06 | No | zinc finger (B-box type) family protein chr2:19615066-19616702 FORWARD Aliases: F17A2 |
| Yes | Yes | 4.93E-05 | No | auxin-responsive AUX/IAA family protein, similar to SP:O24408:AXII_ARATH Auxin-responsiv |
| Yes | Yes | 1.45E-06 | No | expressed protein chr3:6121059-6123017 FORWARD Aliases: MEB5.8 |
| Yes | Yes | 6.17E-05 | No | CCAAT-box binding transcription factor, putative, similar to CAAT-box DNA binding protein s |
| Yes | Yes | 3.84E-06 | No | auxin-responsive protein / indoleacetic acid-induced protein 1 (IAA1), identical to SP:P49677 |
| Yes | Yes | 0.000032 | No | scarecrow-like transcription factor 13 (SCL13) chr4:9660996-9663782 REVERSE Aliases: DL4 |
| Yes | Yes | 8.03E-05 | No | expressed protein, weak similarity to storekeeper protein (Solanum tuberosum) GI:1426847 |
| Yes | Yes | 5.19E-06 | No | WRKY family transcription factor, contains Pfam profile: PF03106 WRKY DNA -binding domai |
| Yes | Yes | 9.59E-06 | No | PHD finger protein-related / SET domain-containing protein (TX4), nearly identical over 285 a |
| Yes | Yes | 4.27E-05 | No | floral homeotic protein APETALA2 (AP2), Identical to (SP:P47927) Floral homeotic protein AP |
| Yes | Yes | 5.77E-05 | No | PHD finger family protein / SET domain-containing protein, contains Pfam domain, PF00628 |
| Yes | Yes | 0.000041 | No | zinc finger (CCCH-type) family protein, contains Pfam domain, PF00642: Zinc finger C-x8-C-x |

| | | | | |
|---|---|---|---|---|
| Yes | Yes | 1.62E-05 | No | transcription factor, putative / E2F-like repressor E2L1 (E2L1), identical to E2F-like repressor |
| Yes | Yes | 1.02E-05 | No | similar to zinc finger (CCCH-type) family protein [Arabidopsis thaliana] (TAIR:At3g06410.1); s |
| Yes | No | #N/A | No | expressed protein chr5:8867843-8869824 REVERSE Aliases: None |
| Yes | Yes | 3.88E-05 | No | expressed protein chr5:17017511-17019835 FORWARD Aliases: MDH9.22, MDH9_22 |
| Yes | Yes | 1.85E-05 | No | similar to glycosyl hydrolase family 14 protein [Arabidopsis thaliana] (TAIR:At2g45880.1); sin |
| Yes | Yes | 1.01E-05 | No | expressed protein, similar to expressed protein [Arabidopsis thaliana] (TAIR:At2g33880.1); si |
| Yes | Yes | 1.45E-05 | Yes | similar to putative zinc finger protein [Oryza sativa (japonica cultivar-group)] (GB:AAT77838. |
| Yes | Yes | 7.34E-07 | Yes | auxin-responsive protein / indoleacetic acid-induced protein 7 (IAA7), identical to SP:Q3882 |
| Yes | Yes | 1.32E-05 | No | homeobox-leucine zipper family protein / homeodomain GLABRA2 like protein 1 (HD-GL2-1) |
| Yes | Yes | 1.22E-05 | No | Dof-type zinc finger domain-containing protein, Zn finger protein BBF2aO -Nicotiana tabacur |
| Yes | Yes | 2.85E-07 | No | zinc finger protein-related, contains Pfam profile PF04396: Protein of unknown function DU |
| Yes | No | #N/A | No | ARID/BRIGHT DNA-binding domain-containing protein, contains Pfam profile PF01388: ARID |
| Yes | Yes | 0.000034 | No | homeobox protein LUMINIDEPENDENS (LD), identical to Homeobox protein LUMINIDEPEND |
| Yes | Yes | 2.29E-05 | No | SET domain-containing protein, contains Pfam profile PF00856: SET domain chr4:8651999- |
| Yes | Yes | 6.94E-07 | No | homeobox-leucine zipper transcription factor (HB-8), identical to HD-zip transcription factor |
| Yes | Yes | 2.96E-07 | No | expressed protein, similar to 6b-interacting protein 1 (NtSIP1) (Nicotiana tabacum) GI:18149 |
| Yes | No | #N/A | No | encodes a member of the DREB subfamily A-5 of ERF/AP2 transcription factor family. The pr |
| Yes | Yes | 4.41E-05 | No | auxin-responsive protein / indoleacetic acid-induced protein 5 (IAA5) / auxin-induced protei |
