## Supplemental Table 6.2 for "A RETINOBLASTOMA-RELATED transcription factor network governs egg cell differentiation and stress response in *Arabidopsis*"

**SI Appendix Table S6.2** List of RBR-regulated transcription factors, and stress and stimulus-responsive transcripts in the egg cell, subtracted from the ovule transcriptomes (B) Stress response genes

| ***Arabidopsis* Locus ID** | **Gene Model Description** | **TF Family Name** |
|---|---|---|
| **I. Enriched in *rbr-3*** | | |
| AT4G25540 | homolog of DNA mismatch repair protein MSH3 | - |
| AT1G14410 | ssDNA-binding transcriptional regulator | Whirly |
| AT5G46210 | cullin4 | - |
| AT1G63940 | monodehydroascorbate reductase 6 | - |
| AT1G13440 | glyceraldehyde-3-phosphate dehydrogenase C2 | - |
| AT1G56450 | 20S proteasome beta subunit G1 | - |
| AT1G09340 | chloroplast RNA binding protein | - |
| AT1G08130 | DNA ligase 1 | - |
| AT2G28190 | copper/zinc superoxide dismutase 2 | - |
| AT3G23830 | glycine-rich RNA-binding protein 4 | - |
| AT3G03300 | dicer-like 2 | - |
| AT1G50500 | Membrane trafficking VPS53 family protein | - |
| AT3G18630 | uracil dna glycosylase | - |
| AT4G37000 | accelerated cell death 2 (ACD2) | - |
| AT2G45640 | SIN3 associated polypeptide P18 | - |
| AT1G74950 | TIFY domain/Divergent CCT motif family protein | ZIM |
| AT5G59880 | actin depolymerizing factor 3 | - |
| AT4G17750 | heat shock factor 1 | HSF |
| AT3G10800 | Basic-leucine zipper (bZIP) transcription factor family protein | bZIP |
| AT5G47390 | myb-like transcription factor family protein | MYB-related |
| AT5G06290 | 2-cysteine peroxiredoxin B | - |
| AT1G44170 | aldehyde dehydrogenase 3H1 | - |
| AT3G56090 | ferritin 3 | - |
| AT1G77310 | wound-responsive family protein | - |
| AT3G12780 | phosphoglycerate kinase 1 | - |
| AT5G54640 | Histone superfamily protein | - |
| AT2G38170 | cation exchanger 1 | - |
| AT5G19330 | ARM repeat protein interacting with ABF2 | - |
| AT5G17310 | UDP-glucose pyrophosphorylase 2 | - |
| AT2G17870 | cold shock domain protein 3 | - |
| AT3G20330 | PYRIMIDINE B | - |
| AT5G59820 | C2H2-type zinc finger family protein | C2H2 |
| AT1G53670 | methionine sulfoxide reductase B 1 | - |
| AT2G25930 | hydroxyproline-rich glycoprotein family protein | - |
| AT2G01440 | DEAD/DEAH box RNA helicase family protein | - |
| AT3G54960 | PDI-like 1-3 | - |
| AT1G29340 | plant U-box 17 | - |
| AT5G49230 | Drought-responsive family protein | - |
| AT5G16970 | alkenal reductase | - |
| AT4G03280 | photosynthetic electron transfer C | - |
| AT5G65020 | annexin 2 | - |
| AT1G65070 | DNA mismatch repair protein MutS%2C type 2 | - |

| | | |
|---|---|---|
| AT3G17210 | heat stable protein 1 | - |
| AT5G59430 | telomeric repeat binding protein 1 | - |
| AT5G05170 | Cellulose synthase family protein | - |
| AT3G10140 | RECA homolog 3 | - |
| AT3G47520 | malate dehydrogenase | - |
| AT3G62980 | F-box/RNI-like superfamily protein | - |
| AT5G55070 | Dihydrolipoamide succinyltransferase | - |
| AT5G47100 | calcineurin B-like protein 9 | - |
| AT1G31470 | Major facilitator superfamily protein | - |
| AT1G66730 | DNA LIGASE 6 | - |
| AT3G24440 | Fibronectin type III domain-containing protein | - |
| AT2G40950 | Basic-leucine zipper (bZIP) transcription factor family protein | bZIP |
| AT1G64790 | ILITYHIA | - |
| AT3G55460 | SC35-like splicing factor 30 | - |
| AT2G30350 | Excinuclease ABC%2C C subunit%2C N-terminal | - |
| AT3G59380 | farnesyltransferase A | - |
| AT5G67590 | NADH-ubiquinone oxidoreductase-like protein | - |
| AT3G52190 | phosphate transporter traffic facilitator1 | - |
| AT4G01370 | MAP kinase 4 | - |
| AT1G49430 | long-chain acyl-CoA synthetase 2 | - |
| AT5G59320 | lipid transfer protein 3 | - |
| AT3G19760 | eukaryotic initiation factor 4A-III | - |
| AT1G22070 | transcription factor TGA3 | bZIP |
| AT1G49950 | telomere repeat binding factor 1 | MYB-related |
| AT1G54100 | aldehyde dehydrogenase 7B4 | - |
| AT1G04400 | cryptochrome 2 | - |
| AT4G00550 | digalactosyl diacylglycerol deficient 2 | - |
| AT3G02140 | AFP2 (ABI five-binding protein 2) family protein | - |
| AT1G65930 | cytosolic NADP+-dependent isocitrate dehydrogenase | - |
| AT2G27050 | ETHYLENE-INSENSITIVE3-like 1 | EIL |
| AT3G22370 | alternative oxidase 1A | - |
| AT4G11600 | glutathione peroxidase 6 | - |
| AT1G66340 | Signal transduction histidine kinase%2C hybrid-type%2C ethylene sensor | - |
| AT3G27820 | monodehydroascorbate reductase 4 | - |
| AT2G17520 | Endoribonuclease/protein kinase IRE1-like protein | - |
| AT5G13850 | nascent polypeptide-associated complex subunit alpha-like protein 3 | - |
| AT1G49300 | RAB GTPase homolog G3E | - |
| AT3G46970 | alpha-glucan phosphorylase 2 | - |
| AT5G52660 | Homeodomain-like superfamily protein | MYB-related |
| AT1G42990 | basic region/leucine zipper motif 60 | bZIP |
| AT3G11930 | Adenine nucleotide alpha hydrolases-like superfamily protein | - |
| AT3G11410 | protein phosphatase 2CA | - |
| AT1G04510 | MOS4-associated complex 3A | - |
| AT1G60490 | vacuolar protein sorting 34 | - |
| AT4G02600 | Seven transmembrane MLO family protein | - |
| AT3G55440 | triosephosphate isomerase | - |
| AT3G06110 | MAPK phosphatase 2 | - |

| | | |
|---|---|---|
| AT2G28290 | P-loop containing nucleoside triphosphate hydrolases superfamily protein | - |
| AT1G09560 | germin-like protein 5 | - |
| AT5G47120 | BAX inhibitor 1 | - |
| AT3G55280 | ribosomal protein L23AB | - |
| AT3G44340 | hypothetical protein | - |
| AT5G53000 | 2A phosphatase associated protein of 46 kD | - |
| AT3G50830 | cold-regulated 413-plasma membrane 2 | - |
| AT2G43520 | trypsin inhibitor protein 2 | - |
| AT4G39640 | gamma-glutamyl transpeptidase 1 | - |
| AT5G54170 | Polyketide cyclase/dehydrase and lipid transport superfamily protein | - |
| AT4G11150 | vacuolar ATP synthase subunit E1 | - |
| AT3G06760 | Drought-responsive family protein | - |
| AT5G21040 | F-box protein 2 | - |
| AT1G66760 | MATE efflux family protein | - |
| AT4G24280 | chloroplast heat shock protein 70-1 | - |
| AT3G23920 | beta-amylase 1 | - |
| AT4G34240 | aldehyde dehydrogenase 3I1 | - |
| AT4G33030 | sulfoquinovosyldiacylglycerol 1 | - |
| AT1G78900 | vacuolar ATP synthase subunit A | - |
| AT4G16950 | Disease resistance protein (TIR-NBS-LRR class) family | - |
| AT2G02860 | sucrose transporter 2 | - |
| AT1G60940 | SNF1-related protein kinase 2.10 | - |
| AT1G53580 | glyoxalase II 3 | - |
| AT2G40300 | ferritin 4 | - |
| AT1G03680 | thioredoxin M-type 1 | - |
| AT3G45140 | lipoxygenase 2 | - |
| AT2G04240 | RING/U-box superfamily protein | - |
| AT2G45790 | phosphomannomutase | - |
| AT5G55390 | ENHANCED DOWNY MILDEW 2 | - |
| AT4G33430 | BRI1-associated receptor kinase | - |
| AT5G04720 | ADR1-like 2 | - |
| | | - |
| **II. depleted in *rbr-3*** | | - |
| AT1G05180 | NAD(P)-binding Rossmann-fold superfamily protein | - |
| AT2G31320 | poly(ADP-ribose) polymerase 2 | - |
| AT1G67500 | recovery protein 3 | - |
| AT1G15020 | quiescin-sulfhydryl oxidase 1 | - |
| AT2G42810 | protein phosphatase 5.2 | - |
| AT1G17880 | basic transcription factor 3 | - |
| AT5G62500 | end binding protein 1B | - |
| AT4G32150 | vesicle-associated membrane protein 711 | - |
| AT1G02500 | S-adenosylmethionine synthetase 1 | - |
| AT2G46370 | Auxin-responsive GH3 family protein | - |
| AT3G10920 | manganese superoxide dismutase 1 | - |
| AT3G08720 | serine/threonine protein kinase 2 | - |
| AT1G14980 | chaperonin 10 | - |
| AT1G33560 | Disease resistance protein (CC-NBS-LRR class) family | - |

| | | |
|---|---|---|
| AT5G14250 | Proteasome component (PCI) domain protein | - |
| AT2G38120 | Transmembrane amino acid transporter family protein | - |
| AT3G59470 | Far-red impaired responsive (FAR1) family protein | - |
| AT3G45640 | mitogen-activated protein kinase 3 | - |
| AT3G11250 | Ribosomal protein L10 family protein | - |
| AT5G08620 | DEA(D/H)-box RNA helicase family protein | - |
| AT1G22280 | phytochrome-associated protein phosphatase type 2C | - |
| AT5G04140 | glutamate synthase 1 | - |
| AT2G26990 | proteasome family protein | - |
| AT3G48750 | cell division control 2 | - |
| AT2G26430 | arginine-rich cyclin 1 | - |
| AT2G25140 | casein lytic proteinase B4 | - |
| AT4G37930 | serine transhydroxymethyltransferase 1 | - |
| AT1G54490 | exoribonuclease 4 | - |
| AT5G12080 | mechanosensitive channel of small conductance-like 10 | - |
| AT3G50820 | photosystem II subunit O-2 | - |
| AT3G06510 | Glycosyl hydrolase superfamily protein | - |
| AT3G23050 | indole-3-acetic acid 7 | AUX-IAA |
| AT3G48190 | Serine/Threonine-kinase ATM-like protein | - |
| AT5G21010 | BTB-POZ and MATH domain 5 | - |
| AT5G50375 | cyclopropyl isomerase | - |
| AT5G63570 | glutamate-1-semialdehyde-2%2C1-aminomutase | - |
| AT4G08500 | MAPK/ERK kinase kinase 1 | - |
| AT1G31812 | acyl-CoA-binding protein 6 | - |
| AT1G77000 | RNI-like superfamily protein | - |
| AT4G32770 | tocopherol cyclase%2C chloroplast / vitamin E deficient 1 (VTE1) / sucrose export defective 1 (SXD1) | - |
| AT3G54720 | Peptidase M28 family protein | - |
| AT4G25050 | acyl carrier protein 4 | - |
| AT1G12860 | basic helix-loop-helix (bHLH) DNA-binding superfamily protein | bHLH |
| AT5G60410 | DNA-binding protein with MIZ/SP-RING zinc finger%2C PHD-finger and SAP domain-containing protein | PHD |
| AT2G19450 | membrane bound O-acyl transferase (MBOAT) family protein | - |
| AT1G49240 | actin 8 | - |
| AT3G18780 | actin 2 | - |
| AT5G65940 | beta-hydroxyisobutyryl-CoA hydrolase 1 | - |
| AT3G12360 | Ankyrin repeat family protein | - |
| AT1G42970 | glyceraldehyde-3-phosphate dehydrogenase B subunit | - |
| AT5G01410 | Aldolase-type TIM barrel family protein | - |
| AT2G27020 | 20S proteasome alpha subunit G1 | - |
| AT2G26890 | DNAJ heat shock N-terminal domain-containing protein | - |

#H3K27M3 target was determined based on normalized read count value present in PlantDHS database for respective gene IDs.

### RBR/E2F binding site was predicted running FIMO algorithm using published E2F binding motifs as templates

| H3K27M3 target | RB/E2F binding site | | TF description |
|---|---|---|---|
| Yes/No | Yes/No | P-value | |
| No | No | #N/A | - |
| Yes | Yes | 9.65E-08 | DNA-binding protein-related, similar to DNA-binding protein p24 GI:9651810 from (Solanum tuberosu |
| No | No | #N/A | - |
| No | No | #N/A | - |
| No | No | #N/A | - |
| No | No | #N/A | - |
| No | No | #N/A | - |
| No | No | #N/A | - |
| No | No | #N/A | - |
| No | No | #N/A | - |
| No | No | #N/A | - |
| No | No | #N/A | - |
| No | No | #N/A | - |
| No | No | #N/A | - |
| No | Yes | 0.0000803 | - |
| Yes | Yes | 0.0000117 | expressed protein chr1:28152236-28154055 REVERSE Aliases: F25A4.8, F25A4_8 |
| No | No | #N/A | - |
| Yes | Yes | 0.000000608 | heat shock factor protein 1 (HSF1) / heat shock transcription factor 1 (HSTF1), identical to heat shock t |
| Yes | Yes | 0.0000487 | bZIP transcription factor family protein, contains Pfam profile: PF00170 bZIP transcription factor; conta |
| Yes | Yes | 0.0000104 | myb family transcription factor, contains Pfam profile: PF00249 myb-like DNA-binding domain chr5:1[illegible] |
| No | No | #N/A | - |
| No | Yes | 0.0000117 | - |
| No | No | #N/A | - |
| No | No | #N/A | - |
| No | No | #N/A | - |
| No | No | #N/A | - |
| No | No | #N/A | - |
| No | No | #N/A | - |
| No | No | #N/A | - |
| No | No | #N/A | - |
| No | No | #N/A | - |
| Yes | Yes | 0.00000358 | zinc finger (C2H2 type) family protein (ZAT12), identical to zinc finger protein ZAT12 (Arabidopsis thalia |
| No | No | #N/A | - |
| No | No | #N/A | - |
| No | No | #N/A | - |
| No | No | #N/A | - |
| No | No | #N/A | - |
| No | No | #N/A | - |
| No | No | #N/A | - |
| No | No | #N/A | - |
| No | No | #N/A | - |
| No | No | #N/A | - |

| | | | |
|---|---|---|---|
| No | No | #N/A | - |
| No | Yes | 0.000062 | - |
| No | Yes | 0.00000166 | - |
| No | No | #N/A | - |
| No | No | #N/A | - |
| No | No | #N/A | - |
| No | No | #N/A | - |
| No | No | #N/A | - |
| No | No | #N/A | - |
| No | No | #N/A | - |
| No | No | #N/A | - |
| Yes | Yes | 0.0000847 | bZIP transcription factor family protein, similar to AtbZIP transcription factor GI:17065880 from (Arabi |
| No | No | #N/A | - |
| No | No | #N/A | - |
| No | No | #N/A | - |
| No | Yes | 1.31E-09 | - |
| No | No | #N/A | - |
| No | No | #N/A | - |
| No | Yes | 0.0000323 | - |
| No | Yes | 0.0000177 | - |
| No | No | #N/A | - |
| No | No | #N/A | - |
| Yes | Yes | 0.0000502 | bZIP family transcription factor (TGA3), identical to transcription factor GI:304113 from (Arabidopsis tl |
| Yes | Yes | 0.0000138 | DNA-binding protein, putative, contains similarity to DNA-binding protein PcMYB1 (Petroselinum crisp |
| No | Yes | 0.0000148 | - |
| No | No | #N/A | - |
| No | No | #N/A | - |
| No | No | #N/A | - |
| No | No | #N/A | - |
| Yes | Yes | 0.0000518 | ethylene-insensitive3-like1 (EIL1), identical to ethylene-insensitive3-like1 GI:2224927 from (Arabidops |
| No | No | #N/A | - |
| No | No | #N/A | - |
| No | No | #N/A | - |
| No | No | #N/A | - |
| No | No | #N/A | - |
| No | No | #N/A | - |
| No | No | #N/A | - |
| No | No | #N/A | - |
| Yes | Yes | 0.00000471 | myb family transcription factor, contains PFAM profile: PF00249 myb-like DNA-binding domain chr5:2 |
| Yes | No | #N/A | AtbZIP60 consists of a bZIP DNA binding domain followed by a putative transmembrane domain. GFP |
| No | No | #N/A | - |
| No | Yes | 0.00000152 | - |
| No | No | #N/A | - |
| No | No | #N/A | - |
| No | No | #N/A | - |
| No | No | #N/A | - |
| No | No | #N/A | - |

| | | | |
|---|---|---|---|
| No | No | #N/A | - |
| No | No | #N/A | - |
| No | No | #N/A | - |
| No | No | #N/A | - |
| No | No | #N/A | - |
| No | No | #N/A | - |
| No | No | #N/A | - |
| No | No | #N/A | - |
| No | No | #N/A | - |
| No | No | #N/A | - |
| No | No | #N/A | - |
| No | No | #N/A | - |
| No | No | #N/A | - |
| No | No | #N/A | - |
| No | No | #N/A | - |
| No | No | #N/A | - |
| No | Yes | 0.000069 | - |
| No | No | #N/A | - |
| No | No | #N/A | - |
| No | No | #N/A | - |
| No | No | #N/A | - |
| No | No | #N/A | - |
| No | No | #N/A | - |
| No | No | #N/A | - |
| No | No | #N/A | - |
| No | Yes | 0.0000173 | - |
| No | Yes | 0.0000698 | - |
| No | No | #N/A | - |
| No | No | #N/A | - |
| No | Yes | 0.0000229 | - |
| No | No | #N/A | - |
| No | | #N/A | - |
| No | | #N/A | - |
| No | Yes | 0.0000393 | - |
| No | No | #N/A | - |
| No | No | #N/A | - |
| No | No | #N/A | - |
| No | No | #N/A | - |
| No | No | #N/A | - |
| No | No | #N/A | - |
| No | No | #N/A | - |
| No | Yes | 0.0000248 | - |
| No | Yes | 0.000017 | - |
| No | No | #N/A | - |
| No | No | #N/A | - |
| No | No | #N/A | - |
| No | No | #N/A | - |

| | | | |
|---|---|---|---|
| No | No | #N/A | - |
| No | Yes | 0.0000155 | - |
| No | No | #N/A | - |
| No | Yes | 0.0000484 | - |
| No | No | #N/A | - |
| No | No | #N/A | - |
| No | No | #N/A | - |
| No | No | #N/A | - |
| No | No | #N/A | - |
| No | No | #N/A | - |
| No | No | #N/A | - |
| No | No | #N/A | - |
| No | No | #N/A | - |
| No | Yes | 0.0000301 | - |
| No | No | #N/A | - |
| No | No | #N/A | - |
| No | No | #N/A | - |
| Yes | Yes | 0.000000734 | auxin-responsive protein / indoleacetic acid-induced protein 7 (IAA7), identical to SP:Q38825:AXI7_AR |
| No | No | #N/A | - |
| No | Yes | 0.0000124 | - |
| No | No | #N/A | - |
| No | No | #N/A | - |
| No | No | #N/A | - |
| No | No | #N/A | - |
| No | No | #N/A | - |
| No | No | #N/A | - |
| No | Yes | 0.0000208 | - |
| No | No | #N/A | - |
| Yes | Yes | 0.0000366 | basic helix-loop-helix (bHLH) family protein / F-box family protein, contains Pfam profiles: PF00646 F-b |
| Yes | Yes | 0.0000145 | similar to putative zinc finger protein [Oryza sativa (japonica cultivar-group)] (GB:AAT77838.1); similar |
| No | Yes | 0.0000162 | - |
| No | No | #N/A | - |
| No | Yes | 0.00000777 | - |
| No | No | #N/A | - |
| No | No | #N/A | - |
| No | No | #N/A | - |
| No | No | #N/A | - |
| No | No | #N/A | - |
| No | No | #N/A | - |
