## Supplemental Table 6.3 for "A RETINOBLASTOMA-RELATED transcription factor network governs egg cell differentiation and stress response in *Arabidopsis*"

**SI Appendix Table S6.3** List of RBR-regulated transcription factors, and stress and stimulus-responsive transcripts in the egg cell, subtracted from the ovule transcriptomes (B) Stress response genes

| *Arabidopsis* Locus ID | Gene Model Description | TF Family Name |
|---|---|---|
| **I. Enriched in *rbr-3*** | | |
| AT3G02140 | AFP2 (ABI five-binding protein 2) family protein | - |
| AT2G19500 | cytokinin oxidase 2 | - |
| AT5G01810 | CBL-interacting protein kinase 15 | - |
| AT1G04710 | peroxisomal 3-ketoacyl-CoA thiolase 4 | - |
| AT1G05710 | basic helix-loop-helix (bHLH) DNA-binding superfamily protein | - |
| AT1G14410 | ssDNA-binding transcriptional regulator | Whirly |
| AT1G17730 | vacuolar protein sorting 46.1 | - |
| AT1G22070 | transcription factor TGA3 | bZIP |
| AT1G49040 | stomatal cytokinesis defective / SCD1 protein (SCD1) | - |
| AT1G66340 | Signal transduction histidine kinase%2C hybrid-type%2C ethylene sensor | - |
| AT2G04240 | RING/U-box superfamily protein | - |
| AT2G13540 | ARM repeat superfamily protein | - |
| AT2G26070 | protein REVERSION-TO-ETHYLENE SENSITIVITY-like protein (DUF778) | - |
| AT2G27050 | ETHYLENE-INSENSITIVE3-like 1 | EIL |
| AT2G45640 | SIN3 associated polypeptide P18 | - |
| AT3G11410 | protein phosphatase 2CA | - |
| AT3G27310 | plant UBX domain-containing protein 1 | - |
| AT3G45140 | lipoxygenase 2 | - |
| AT3G59380 | farnesyltransferase A | - |
| AT4G01370 | MAP kinase 4 | - |
| AT4G03560 | two-pore channel 1 | - |
| AT4G33430 | BRI1-associated receptor kinase | - |
| AT4G37770 | 1-amino-cyclopropane-1-carboxylate synthase 8 | - |
| AT4G39850 | peroxisomal ABC transporter 1 | - |
| AT5G13300 | ARF GTPase-activating protein | - |
| AT5G25900 | GA requiring 3 | - |
| AT5G47100 | calcineurin B-like protein 9 | - |
| AT5G54500 | flavodoxin-like quinone reductase 1 | - |
| AT5G59430 | telomeric repeat binding protein 1 | - |
| AT5G64813 | Ras-related small GTP-binding family protein | - |
| AT1G13980 | sec7 domain-containing protein | - |
| AT1G44170 | aldehyde dehydrogenase 3H1 | - |
| AT1G49950 | telomere repeat binding factor 1 | MYB-related |
| AT1G54100 | aldehyde dehydrogenase 7B4 | - |
| AT1G74950 | TIFY domain/Divergent CCT motif family protein | ZIM |
| AT2G17800 | RAC-like 1 | - |
| AT3G58680 | multiprotein bridging factor 1B | MBF1 |
| AT4G25000 | alpha-amylase-like protein | - |
| AT5G59320 | lipid transfer protein 3 | - |
| AT1G33410 | SUPPRESSOR OF AUXIN RESISTANCE1 | - |
| AT1G54270 | eif4a-2 | - |
| AT2G06850 | xyloglucan endotransglucosylase/hydrolase 4 | - |

| | | |
|---|---|---|
| AT2G29420 | glutathione S-transferase tau 7 | - |
| AT5G02310 | proteolysis 6 | - |
| AT5G20990 | molybdopterin biosynthesis CNX1 protein / molybdenum cofactor biosynthesis enzyme CNX1 (CNX1) | - |
| AT5G26780 | serine hydroxymethyltransferase 2 | - |
| AT5G52660 | Homeodomain-like superfamily protein | MYB-related |
| AT4G26200 | 1-amino-cyclopropane-1-carboxylate synthase 7 | - |
| AT4G34240 | aldehyde dehydrogenase 3I1 | - |
| AT5G05170 | Cellulose synthase family protein | - |
| AT5G19330 | ARM repeat protein interacting with ABF2 | - |
| AT5G47390 | myb-like transcription factor family protein | MYB-related |
| AT3G62980 | F-box/RNI-like superfamily protein | - |
| AT1G49430 | long-chain acyl-CoA synthetase 2 | - |
| AT2G25930 | hydroxyproline-rich glycoprotein family protein | - |
| AT2G30980 | SHAGGY-related protein kinase dZeta | - |
| AT5G01810 | CBL-interacting protein kinase 15 | - |

#H3K27M3 target was determined based on normalized read count value present in PlantDHS database for respective gene IDs.

### RBR/E2F binding site was predicted running FIMO algorithm using published E2F binding motifs as templates

| H3K27M3 target | RB/E2F binding site | | stress responsive |
|---|---|---|---|
| Yes/No | Yes/No | P-value | |
| No | No | #N/A | Yes |
| No | Yes | 0.00000854 | No |
| No | Yes | 0.0000382 | No |
| No | Yes | 0.0000539 | No |
| No | Yes | 0.00000879 | No |
| Yes | Yes | 9.65E-08 | Yes |
| No | Yes | 0.00000272 | No |
| Yes | Yes | 0.0000502 | Yes |
| No | Yes | 0.0000739 | No |
| No | No | #N/A | Yes |
| No | Yes | 0.0000698 | Yes |
| No | Yes | 0.0000073 | No |
| No | Yes | 0.0000564 | No |
| Yes | Yes | 0.0000518 | Yes |
| No | Yes | 0.0000803 | Yes |
| No | Yes | 0.00000152 | Yes |
| No | Yes | 0.0000198 | No |
| No | Yes | 0.0000173 | Yes |
| No | Yes | 1.31E-09 | Yes |
| No | Yes | 0.0000323 | Yes |
| No | Yes | 0.0000157 | No |
| No | Yes | 0.0000229 | Yes |
| No | Yes | 0.0000956 | No |
| No | Yes | 0.0000756 | No |
| No | Yes | 0.0000222 | No |
| No | Yes | 0.0000158 | No |
| No | No | #N/A | Yes |
| No | Yes | 0.0000156 | No |
| No | Yes | 0.000062 | Yes |
| No | Yes | 0.0000427 | No |
| No | Yes | 0.00000041 | No |
| No | Yes | 0.0000117 | Yes |
| Yes | Yes | 0.0000138 | Yes |
| No | Yes | 0.0000148 | Yes |
| Yes | Yes | 0.0000117 | Yes |
| No | Yes | 0.0000268 | No |
| Yes | Yes | 0.0000414 | No |
| No | Yes | 0.0000415 | No |
| No | No | #N/A | Yes |
| No | Yes | 0.0000441 | No |
| No | Yes | 0.000076 | No |
| No | Yes | 0.00000915 | No |

| | | | |
|---|---|---|---|
| No | Yes | 0.00000962 | No |
| No | Yes | 0.0000577 | No |
| No | Yes | 0.0000124 | No |
| No | Yes | 0.0000768 | No |
| Yes | Yes | 0.00000471 | Yes |
| No | Yes | 0.0000767 | No |
| No | Yes | 0.000069 | Yes |
| No | Yes | 0.00000166 | Yes |
| No | No | #N/A | Yes |
| Yes | Yes | 0.0000104 | Yes |
| No | No | #N/A | Yes |
| No | Yes | 0.0000177 | Yes |
| No | No | #N/A | Yes |
| No | Yes | 0.00000671 | No |
| No | Yes | 0.0000382 | No |

| TF description |
| --- |
| - |
| - |
| - |
| - |
| - |
| DNA-binding protein-related, similar to DNA-binding protein p24 GI:9651810 from (Solanum tuberosum) \| chr1:4929118-4930888 REVERSE \| Aliases: F14L17 |
| - |
| bZIP family transcription factor (TGA3), identical to transcription factor GI:304113 from (Arabidopsis thaliana) \| chr1:7789497-7792103 FORWARD \| Aliases: F |
| - |
| - |
| - |
| - |
| - |
| ethylene-insensitive3-like1 (EIL1), identical to ethylene-insensitive3-like1 GI:2224927 from (Arabidopsis thaliana) \| chr2:11552873-11555371 FORWARD \| Ali |
| - |
| - |
| - |
| - |
| - |
| - |
| - |
| - |
| - |
| - |
| - |
| - |
| - |
| - |
| - |
| - |
| - |
| - |
| DNA-binding protein, putative, contains similarity to DNA-binding protein PcMYB1 (Petroselinum crispum) gi:2224899:gb:AAB61699 \| chr1:18497832-18500 |
| - |
| expressed protein \| chr1:28152236-28154055 REVERSE \| Aliases: F25A4.8, F25A4_8 |
| - |
| ethylene-responsive transcriptional coactivator, putative, similar to ethylene-responsive transcriptional coactivator (Lycopersicon esculentum) gi:5669634:gb: |
| - |
| - |
| - |
| - |
| - |

-
-
-
-
myb family transcription factor, contains PFAM profile: PF00249 myb-like DNA-binding domain | chr5:21376285-21379384 REVERSE | Aliases: F6N7.15, F6N7
-
-
-
-
myb family transcription factor, contains Pfam profile: PF00249 myb-like DNA-binding domain | chr5:19244017-19246085 FORWARD | Aliases: MQL5.25, MC
-
-
-
-
-
