## Supplemental Table 1,2,3,4, and all supplemental figure for "A RETINOBLASTOMA-RELATED transcription factor network governs egg cell differentiation and stress response in *Arabidopsis*"

### A transcription factor network impinges on eggs (short title)

### Supporting Information

**Table S1. A hemizygous transgene *GFP-gRBR* fully restores fertility of *rbr-3* gametophytes in *rbr-3/RBR; GFP-gRBR*$^{h}$.**

| | viable seeds | aborted ovules | total seed count | ***P*** **value (Fisher's exact test)** |
|---|---|---|---|---|
| expected frequency† | 0.75 | 0.25 | 1 | |
| expected phenotypic ratio | 453 | 151 | 604 | |
| observed phenotypic ratio | 440 | 164 | 604 | 0.4317 $^{ns}$ |

$^{h}$ hemizygous

† expected frequency is calculated based on abortion of *rbr-3* ovules [22] and full viability of *rbr-3; GFP-gRBR* female gametophytes

$^{ns}$not significant difference: Note that there are no significant deviations between the expected and observed ratio.

**Table S2. Progeny test confirms 'two independent loci' complementation of *rbr-3* allele with *GFP-gRBR* in *rbr-3/RBR; GFP-gRBR*$^{h}$ background. Note that offspring was scored based on antibiotic resistance of *rbr-3* T-DNA.**

| | resistant seedlings | sensitive seedlings | total seedling count | ***P*** **value (Fisher's exact test)** |
|---|---|---|---|---|
| expected frequency† | 0.569 | 0.431 | 1 | |
| expected phenotypic ratio | 159 | 120 | 279 | |
| observed phenotypic ratio | 158 | 121 | 279 | 1.0000 $^{ns}$ |

$^{h}$ hemizygous

† expected frequency is calculated based on absence of *rbr-3* transmission to the progeny through the female and 10% transmission via male gametes [22] and full rescue of *rbr-3 by GFP-gRBR* transgene

$^{ns}$not significant difference: Note that there are no significant deviations between the expected and observed ratio.

**Table S3. A hemizygous transgene *pEC-gRBR* partially restores fertility of *rbr-3* gametophytes in *rbr-3/RBR; pEC1-gRBR*$^{h}$ background.**

| | viable seeds | aborted ovules | total seed count | ***P*** **value (Fisher's exact test)** |
|---|---|---|---|---|
| expected frequency† | 0.75 | 0.25 | 1 | |
| expected phenotypic ratio | 80 | 27 | 107 | |
| observed phenotypic ratio in *rbr-3/RBR* | 357 | 299 | 656 | |
| observed phenotypic ratio vs expected | 72 | 35 | 107 | 0.2915 $^{ns}$ |
| observed phenotypic ratio vs *rbr-3* | 72 | 35 | 107 | 0.0154* |

$^{h}$ hemizygous

† expected frequency is calculated based on abortion of *rbr-3* ovules [22] and full viability of *rbr-3; pEC-gRBR* female gametophytes

$^{ns}$not significant difference: Note that there are no significant deviations between the expected rescue seed set and observed ratio.

* statistically significant difference: Note that there are significant deviations between seed set of *rbr-3* alone and observed ratio in *rbr-3/RBR* in presence of *pEC- gRBR*.

**Table S4. Progeny test confirms partial 'two independent loci' complementation of *rbr-3* allele with *pEC-gRBR* in *rbr-3/RBR; pEC-gRBR*$^{h}$ background. Note that offspring was scored based on antibiotic resistance of *rbr-3* T-DNA.**

| | resistant seedlings | sensitive seedlings | total seedling count | ***P*** **value (Fisher's exact test)** |
|---|---|---|---|---|
| expected frequency† | 0.569 | 0.431 | 1 | |
| expected phenotypic ratio | 103 | 78 | 181 | |
| observed phenotypic ration in *rbr-3/RBR* | 35 | 314 | 349 | |
| observed phenotypic ratio vs expected | 44 | 137 | 181 | 0.0001* |
| observed phenotypic ratio vs *rbr-3/RBR* | 44 | 137 | 181 | 0.0001* |

$^{h}$ hemizygous

† expected frequency is calculated based on absence of *rbr-3* transmission to the progeny through the female and 10% transmission via male gametes [22] and full rescue of *rbr-3 by pEC-gRBR* transgene

* statistically significant difference: Note that there are significant deviations between the expected and observed ratio as well as from observed ratio for *rbr-3/RBR* alone.

**Table S5. Previously validated egg cell expressed transcripts showing deregulation in *rbr-3* ovule transcriptome**

**Table S6. List of RBR-regulated transcription factors, and stress and stimulus-responsive transcripts in the egg cell, subtracted from the ovule transcriptomes**

**Table S7. List of egg cell-expressed RBR interactors used for building protein interaction network.**

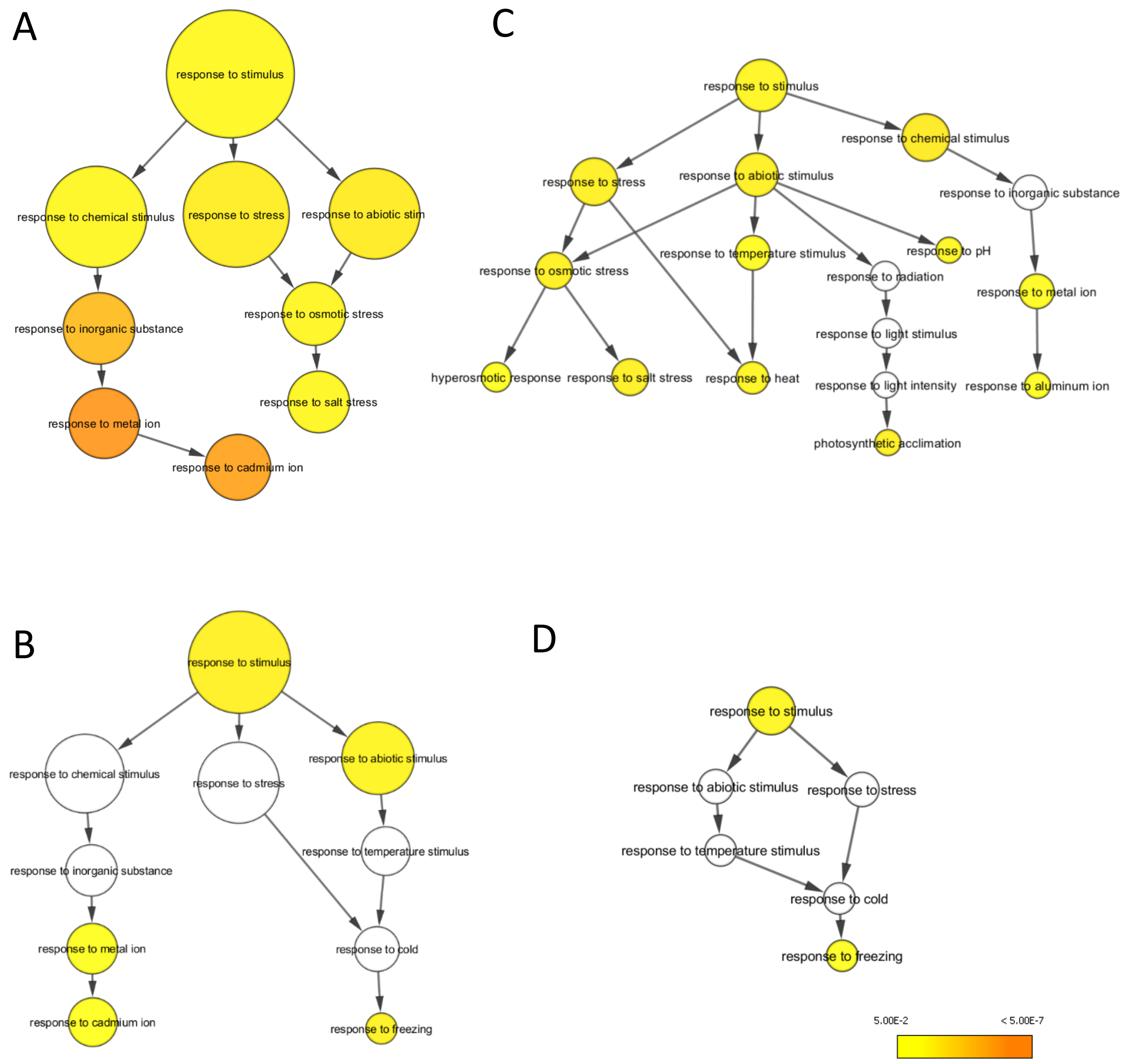


**Fig. S1. Overall gene ontology enrichment of stress and stimulus responsive genes and transcription factors in *rbr-3* egg cells.** Gene ontology clustering for abiotic stress and stimulus enriched (**A**) or depleted (**B**) in *rbr-3* ovules, and enriched (**C**) or depleted (**D**) in *rbr-3* egg cells.

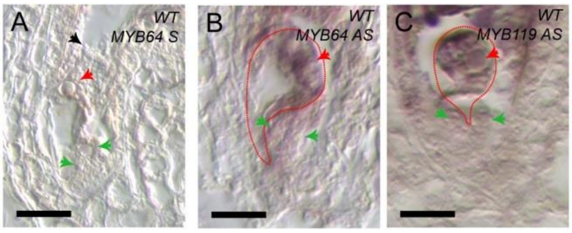


**Fig. S2. *MYB64* and *MYB119* transcripts in the mature embryo sac.** mRNA *in situ* hybridization: (**A**) Sense (*S*) probes shows no signal, while anti-sense probes (*AS*) detect (**B**) *MYB64* and (**C**) *MYB119* mRNA in the mature embryo sac, and specifically in the egg cell. Scale bar=20µm.

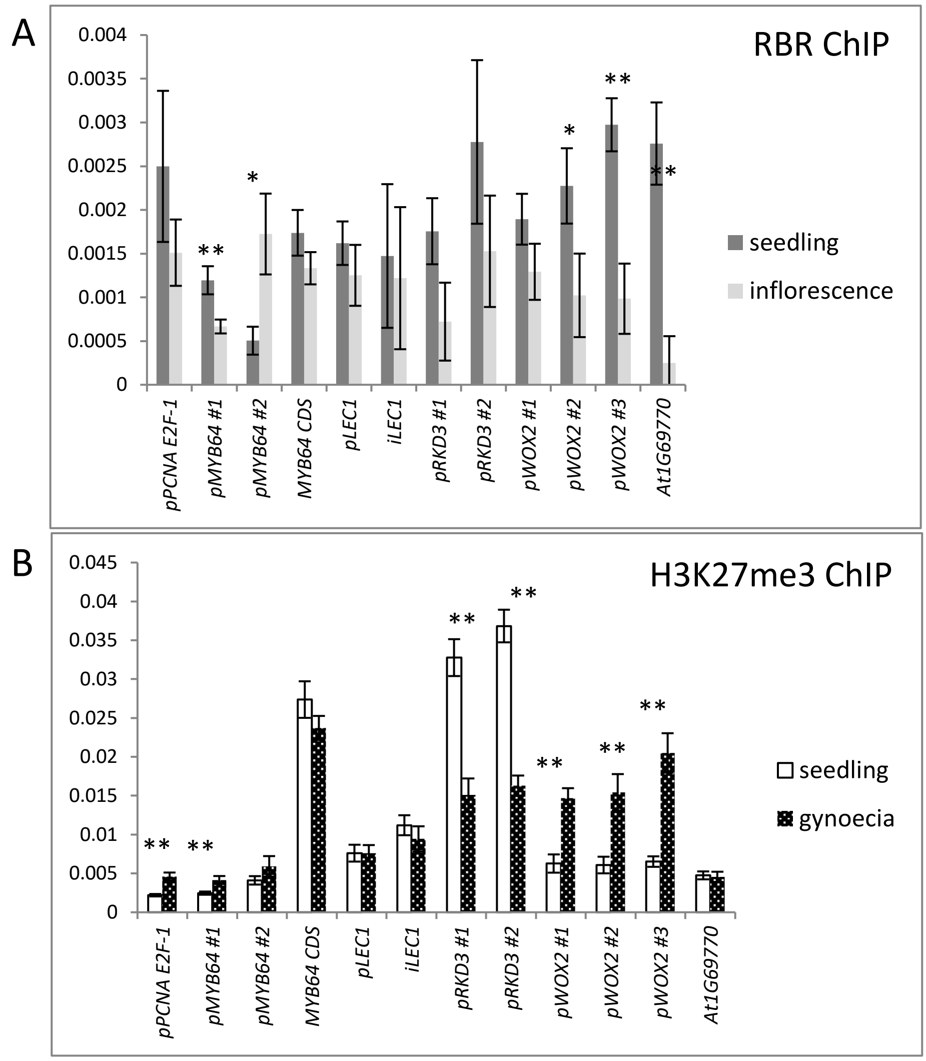


**Fig. S3. Chromatin Immunoprecipitation in reproductive tissues in comparison to vegetative stages in *Arabidopsis.*** (**A**) ChIP for RBR or (**B**) for the PRC2-specific H3K27me3 binding. Relative real-time qPCR data normalized by input. *PCNA* locus was used as a positive control for RBR binding but negative for H3K27me3. Significant difference is indicated between seedling and inflorescence tissues: **α ≤ 0.01; *α ≤ 0.05.

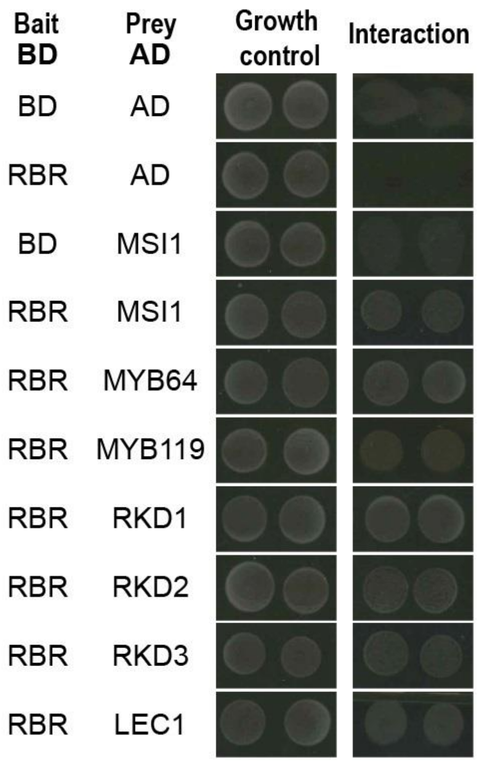


**Fig. S4. RBR interacts with egg-cell expressed transcription factors.** RBR protein-protein interactions identified by LexA-based yeast-two-hybrid assay.

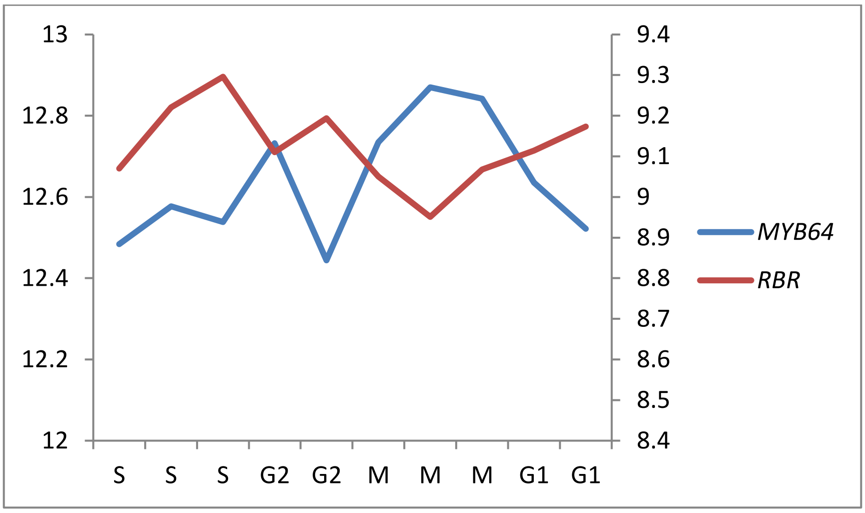


**Fig. S5. Cell-cycle-dependent expression of *MYB64*.** *MYB64* transcript signals change with cell cycle progression in synchronized *Arabidopsis* cell culture in an opposite manner to *RBR* [46].

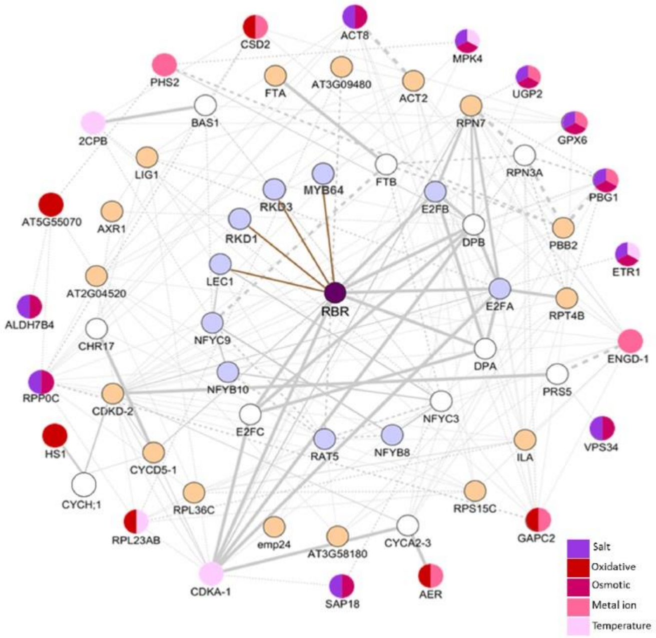


**Fig. S6. RBR connects transcriptional regulation and stress response shared by PRC2.** A protein interaction network connecting RBR, cell cycle, transcription factors and stress-responsive genes. Transcription factors enriched in *rbr-3* egg cell transcriptome connect to stress responsive
